# Dual-Lineage Human 3D Bone Niche Model Reveals Osteoclast-Driven Osteomimicry in Prostate Cancer

**DOI:** 10.1101/2025.11.18.689049

**Authors:** Andrea Mazzoleni, Robin Dolgos, Thomas Menter, Boris Dasen, Arnaud Scherberich, Clémentine Le Magnen, Manuele G Muraro, Ivan Martin

## Abstract

Bone is the predominant site of metastasis in advanced prostate cancer (PCa), yet the mechanisms governing tumor-bone interactions remain incompletely understood. A modular human three-dimensional (3D) in vitro bone niche model is developed that integrates osteoblasts and osteoclasts within a mineralized scaffold, recreating an endosteal-like microenvironment for co-culture with PCa cell lines and patient-derived organoids (PDOs). The engineered construct maintains osteoblastic differentiation and supports osteoclastogenesis, confirmed by lineage markers including osteocalcin, osteopontin (OPN), and tartrate-resistant acid phosphatase (TRAP). Co-culture with PCa cells downregulates osteoblast- and osteoclast-associated genes (IBSP, OPN, TRAP) in bone cells, suggesting tumor-mediated suppression of bone remodeling. Conversely, co-cultured PCa cells exhibit niche-dependent osteomimicry, characterized by upregulation of osteoblastic (SPARC, BGLAP) and osteoclastic (TRAP) markers and strongly regulated by the presence of osteoclasts. The platform also supports engraftment and proliferation of PDOs without PCa-specific exogenous growth factors, underscoring its translational relevance. This osteoblastic-osteoclastic niche model thus provides a human system capturing PCa-bone cell interactions, with potential utility to investigate therapeutic responses in a clinically relevant context.

**Table of Contents:** A human 3D bone niche integrating osteoblasts and osteoclasts enables co-culture with prostate cancer cell lines and patient-derived organoids. The engineered niche (i) models reciprocal phenotypic regulation between bone and cancer cells, (ii) captures osteoclast-enhanced osteomimicry in tumor cells and (iii) establishes a biomimetic platform for mechanistic studies of metastatic prostate cancer cells in bone.

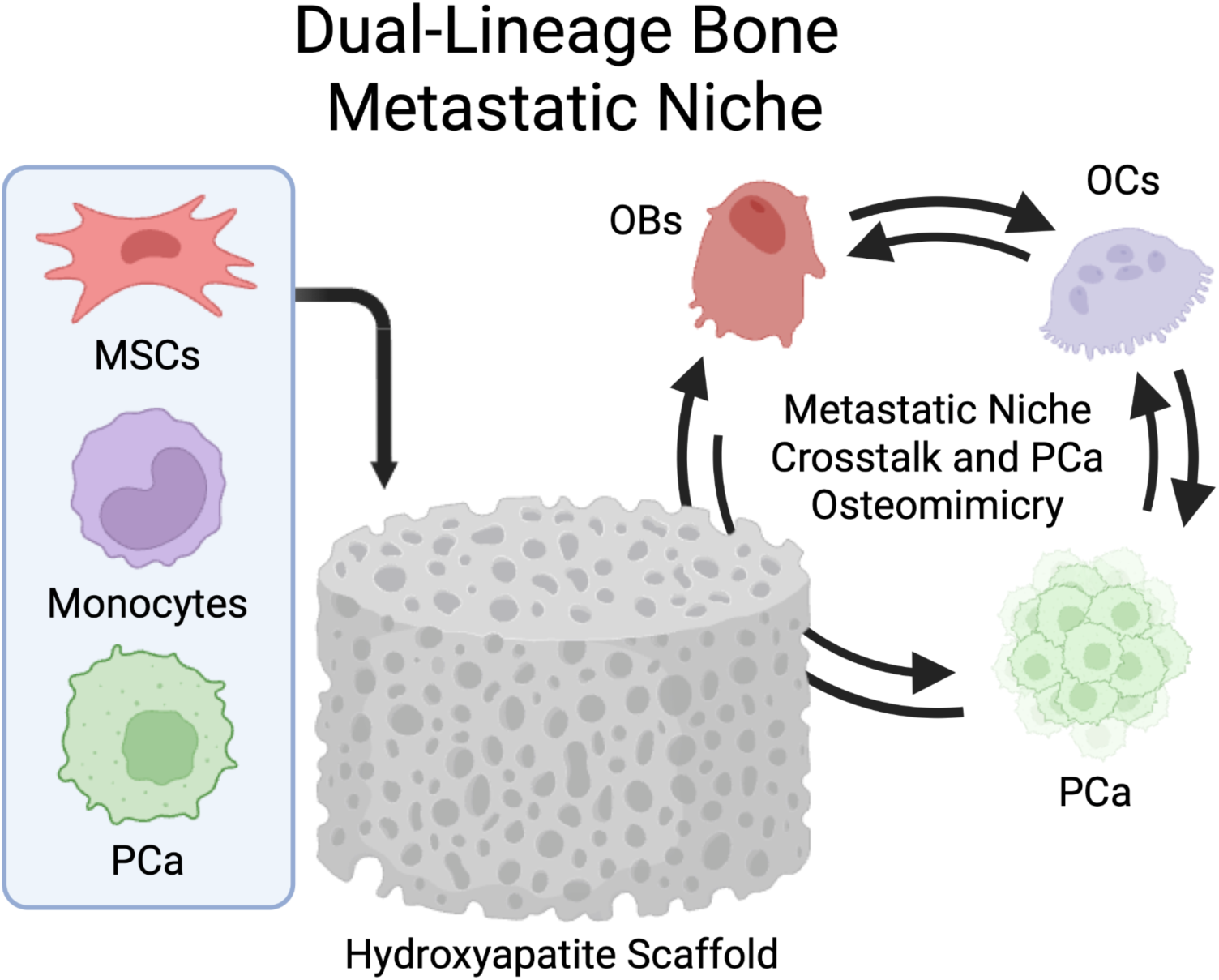

## 1. Introduction

Prostate cancer (PCa) is one of the most prevalent malignancies in men, accounting for 20% of new cancer diagnoses and representing the third leading cause of cancer-related death among men in Europe [1]. While localized PCa is often curable, a significant portion of patients eventually progress to distant metastases, with bone being the most common site (82%) [2]. Once established in bone, metastasis-related events profoundly compromise patient survival and quality of life, underscoring the urgent need to understand better the crosstalk between PCa cells and the bone microenvironment. This specialized niche actively supports metastatic PCa (mPCa) growth and expansion through reciprocal interactions between tumor cells and resident bone cells, disrupting the balance between osteoblastic bone formation and osteoclastic bone resorption [3]. Despite extensive research, the mechanisms and molecular mediators governing these processes remain not fully understood. Advancing our understanding of PCa-bone interactions may pave the way for more effective or novel approaches to manage late-stage mPCa [4].

To dissect cellular interactions in the context of mPCa, several models have been developed, predominantly relying on animal-based systems using cell line- and patient-derived xenografts, transgenic mice, or bone implant models [5]. While these systems have significantly advanced our understanding of the biology of established bone metastases, fundamental physiological differences between species limit fidelity to clinical scenarios. This disconnect is reflected in the low translational success rate of preclinical findings from animal models when tested in human clinical trials [6]. As a result, there is a pressing need for more predictive, ethically-responsible, and 3R-compliant platforms.

*In vitro* 3D metastatic models based on human cells have emerged as a powerful alternative, offering enhanced biomimicry through physiologically relevant cell-matrix interactions while enabling modular co-culture of stromal, immune, and cancer cell types in controlled microenvironments [7,8]. These models hold great promise for improving mechanistic insights and developing therapeutic strategies in the context of bone metastatic PCa [9–11]. However, probably due to the lack of important cellular components of the bone niche, existing models cannot capture pathological processes which may be critical to drive persistence of PCa cells into bone through adaptation and acquisition of bone-like characteristics, e.g. the phenomenon known as osteomimicry [12].

Historically, most PCa bone metastasis models have focused on the interaction between PCa cells and osteoblasts, reflecting the predominance of osteoblastic lesions and pathological bone formation observed in mPCa patients [13–16]. However, this osteoblast-centric approach fails to capture the reciprocal contributions of osteoclasts, whose activity not only mediates bone resorption but also fuels osteoblast activation and tumor growth. In fact, osteoclasts play essential roles in early metastatic seeding by releasing matrix-derived growth factors and modulating osteoblast activity through paracrine signaling [17]. Excluding osteoclasts, therefore, oversimplifies the complex dynamics of the metastatic bone niche and limits mechanistic insight into the vicious cycle of remodeling [18]. When osteoclasts are included, their differentiation is often induced by high doses of exogenous RANKL, which bypasses the endogenous osteoblast–osteoclast signaling axis and may distort the niche’s biology [19]. In addition to cellular interactions, the mineralized extracellular matrix is a defining feature of bone that dynamically regulates osteoblast and osteoclast activity. Its mechanical properties, particularly rigidity and mineral content, promote osteoblast differentiation and function while also modulating osteoclast adhesion, polarization, and resorptive activity, thereby shaping the dynamics of the metastatic niche [20,21]. A mineralized, rigid surface is thus critical to provide physiologically relevant cues, which explains why many previous 3D models of PCa-bone interactions have relied on co-cultures with ex vivo bone tissue from mice or patients [22–24].

Here, we developed a fully human, engineered *in vitro* 3D model of the metastatic bone niche to test the hypothesis that osteoclasts play a critical role in shaping tumor-bone crosstalk. Building on established differentiation protocols [25–27], the platform integrates both osteoblastic and osteoclastic precursors within a mineralized scaffold and supports controlled co-culture with PCa cell lines and a patient-derived organoid-to-xenograft-to-organoid model (PDOXO). By enabling osteoclastogenesis through osteoblast-derived cues rather than exogenous RANKL, the system overcomes a significant limitation of osteoblast-centric systems.

## 2. Results

### 2.1. Engineered niches recapitulate osteoblastic and osteoclastic features of the bone microenvironment

To engineer dual-lineage osteoblastic-osteoclastic niches (OBCNs), we cultured human BM-MSCs (bone marrow derived mesenchymal stromal cells) from different donors (total of 3) on hydroxyapatite-based scaffolds for a total of 4 weeks under conditions promoting osteoblastic differentiation, followed by loading osteoclast progenitors and culturing the constructs in osteoclastic medium (OCM) for an additional 2 weeks (Figure 1A). Osteoblastic-only niches (OBNs) were established by omitting the osteoclastic precursors.

We first evaluated osteoblastic differentiation by gene expression analysis (qRT-PCR; Figure 1B, S. Figure 1A-1D). Alkaline phosphatase (ALPL), a key enzyme in matrix mineralization, showed comparable expression in OBCNs and OBNs across donors and timepoints. BGLAP, encoding osteocalcin, a late osteoblast marker, was consistently downregulated in OBCNs, with decreases ranging from 3-fold to 11-fold at TP1 (time point 1, 7 days after osteoclast seeding; p < 0.05) and further reduced at TP2 (timepoint 2, 14 days after osteoclast seeding; up to 9-fold; p < 0.05), as expected by the absence of osteogenic stimuli. Expression of bone morphogenetic protein 2 (BMP2), a hallmark for osteoblast differentiation and bone formation, remained largely stable among donors and time points (mean < 2-fold). In contrast, integrin-binding sialoprotein (IBSP), involved in matrix mineralization, showed strong donor- and timepoint-dependent variation: IBSP was downregulated in OBCNs from BM293 at both timepoints (p < 0.05), but upregulated in BM267 and BM256 at TP1 (Figure 1B, S. Figure 1A). Overall, gene expression showed differential expression of key markers of osteoblastic commitment, maintaining ALPL and BMP2 expression in OBCNs while downregulating BGLAP and showing donor-specific variation for IBSP. Immunohistochemistry (IHC) analysis revealed osteocalcin-positive cells and matrix-associated IBSP across all conditions. Fusiform cell morphology, typical of early-stage osteoblasts and osteoblastic progenitors, as well as the presence of osteopontin-positive osteoblasts lining the mineral-rich areas of the scaffolds, was evident in H&E-stained sections both confirmed by a pathologist (Figure 1C, S. Figure 2A–2C). Thus, our imaging findings overall confirm maintenance of osteoblastic commitment in both OBCNs and OBNs.

To evaluate osteoclastic differentiation, we examined the expression of tartrate-resistant acid phosphatase (TRAP), a canonical osteoclast marker, alongside osteopontin (OPN), a matrix-associated protein expressed by both osteoblasts and active osteoclasts. OPN was consistently and significantly upregulated in OBCNs across all donors and timepoints, with an increase ranging from 41-fold to 982-fold (Figure 1B, and S. Figure 1A-1D). TRAP, a canonical osteoclast marker, showed similarly robust induction, with an increase exceeding 264-fold in all OBCN donors (Figure 1B, S. Figure 1A-1D). TRAP-positive multinucleated cells were observed exclusively in OBCNs via IHC staining (Figure 1C, S. Figure 2A–2C).

Together, these findings demonstrate that the OBCN platform supports the differentiation and coexistence of both osteoblasts and osteoclasts, and indicate that the presence of osteoclasts modulate the transcriptional profile of osteoblastic cells. Overall, the OBCN recapitulates phenotypic features of the bone microenvironment.

**Figure 1.**
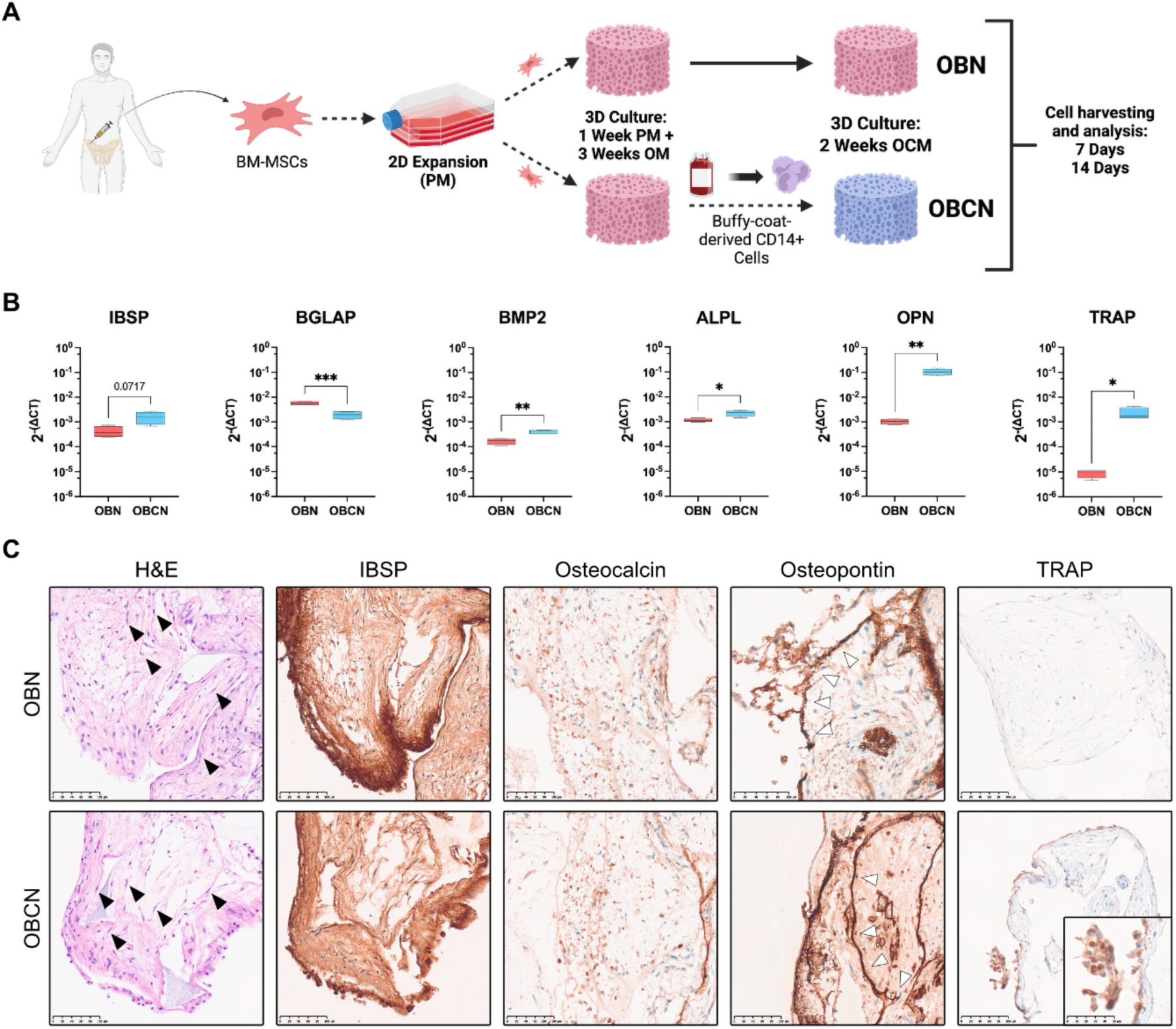
Engineered dual-lineage human 3D bone niches integrating osteoblastic and osteoclastic cells. (A) Schematic representation of the methodology employed to engineer the niches and analyze them. Figure created with BioRender.com. (B) qRT-PCR analysis of osteoblastic and osteoclastic markers IBSP, BGLAP, BMP2, ALPL, OPN, and TRAP in OBNs (red) and OBCNs (blue) on the mixed osteoblast and osteoclast population. Expression level was normalised to GAPDH and calculated using the 2^-ΔCT^ method. Each graph represents data from a singular donor, BM267, at the first analysis timepoint (TP1), with up to n=4 experimental replicates. Comparisons were done via Welch t-test, with statistically significant changes being shown in the graph (✱: p < 0.05; ✱✱: p < 0.005; ✱✱✱: p < 0.0005). (C) Immunohistochemistry imaging of OBNs and OBCNs, comparing H&E staining and protein level expression of IBSP, osteocalcin, osteopontin, and TRAP (Scale bar: 100 µm). Black arrowheads in H&E images indicate cells with fusiform morphology, indicative of osteoblasts. White arrowheads in osteopontin images indicate areas adjacent to scaffold-derived hydroxyapatite. Detailed insets in TRAP images highlight multi-nucleated TRAP+ cells (Scale bar: 50 µm). The data are related to donor BM267 at TP1. Other donors and timepoints are shown in S. Figure 2.

### 2.2. OBCNs support prostate cancer cell engraftment

To evaluate whether the engineered niches support tumor cell engraftment, we seeded fluorescently labeled LNCaP (mEmerald) and PC3 (mCherry) cells into both OBNs and OBCNs, hereafter referred to as L-OBN/OBCN and P-OBN/OBCN, respectively. These two cell lines differ markedly in molecular features: LNCaP cells are androgen receptor–positive (AR⁺) and retain epithelial characteristics, while PC3 cells are AR⁻, display more mesenchymal-like features, and represent an aggressive, castration-resistant tumor phenotype. Recovered tumor cells were quantified via FACS and used for downstream analysis (Figure 2A).

A modestly higher percentage of PC3 cells was recovered from P-OBCNs than from LNCaP cells in L-OBCNs at both timepoints and donors (Figure 2B). This likely reflects intrinsic differences in growth and adaptation between the two cell lines, as PC3 cells exhibit a more aggressive, androgen-independent phenotype. Engraftment of epithelial tumor cells in the niches was confirmed by H&E staining and further validated by E-Cadherin immunostaining. LNCaP and PC3 cells displayed distinct phenotypes and spatial growth patterns within the niches. PC3 cells appeared more flattened and dispersed, with irregular borders and loose cell-cell contacts, while LNCaP cells formed compact, rounded clusters. These morphological differences were accompanied by a weaker E-Cadherin signal in PC3 cells, consistent with their mesenchymal-like features (Figure 2C). Whole-mount immunofluorescence further revealed that LNCaP cells organized into larger, compact 3D aggregates, whereas PC3 cells were more diffusely distributed throughout the construct, reflecting their divergent modes of tumor growth in a bone-like microenvironment. These observations were consistent across donors and timepoints (Figure 2C, S. Figure 3A–4C). Additional evidence of PCa cell engraftment was provided by scanning electron microscopy, which indicated co-localization of epithelial and osteoclast precursors (S. Figure 4).

Together, these findings demonstrate that OBCNs robustly support PCa cell engraftment.

**Figure 2.**
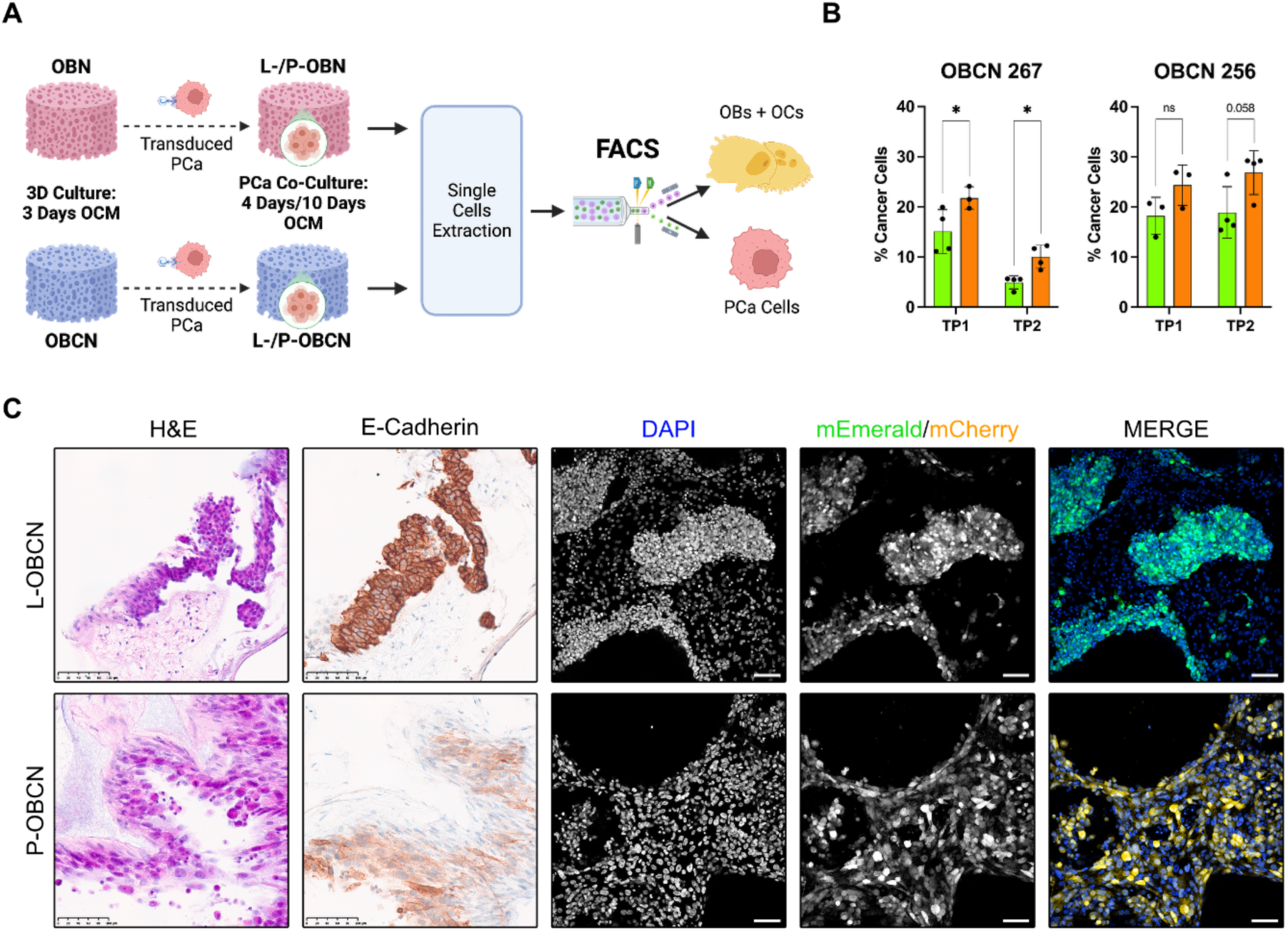
FACS and imaging reveal successful engraftment of PCa cell lines in OBCNs. (A) Schematic representation of the culture process for PCa co-cultured niches and the procedure used for separation of osteoblasts and osteoclasts from PCa cells residing in the niches using FACS. Figure created with BioRender.com. (B) Percentage of the PCa cells retrieved from sorting of OBCNs 267 and 256 is shown for both LNCaP and PC3 lines (up to n = 4 samples per niche type) in green and orange, respectively. The difference between LNCaP and PC3 cells extracted from niches was statistically significant for OBCN 267 (p < 0.05, ✱ per Welch t-test). (C) Representative examples of IHC and IF imaging of L-OBCN and P-OBCN niches, via H&E staining, E-Cadherin staining, and the fluorescent reporters mEmerald and mCherry (Scale bar: 100 µm). Data related to donor BM267 at TP1. Other donors and timepoints are shown in S. Figure 3, 4, and 5.

### 2.3. PCa cell lines differentially downregulate osteoclastic gene expression in OBCNs

Having established the successful engraftment of PCa cells within the bone-like niches, we next addressed whether PCa cells modulated gene and protein expression profiles of the OBCN niche upon co-culture, using qRT-PCR on sorted niche cells and IHC on intact constructs.

Histological analysis revealed no overt differences in protein expression patterns of bone-related markers between OBCNs co-cultured with PCa cells (L-OBCNs and P-OBCNs) and cancer cell-free OBCNs (Figure 3A, S. Figure 5). IBSP retained its matrix-associated localization, while osteocalcin (OCN) was expressed in osteoblastic cells throughout the niche. Osteopontin (OPN) remained strongly expressed in cells lining hydroxyapatite-rich areas, and TRAP-positive multinucleated cells were visible across all conditions. Osteonectin (SPARC), a promoter of osteoblast function and modulator of mineralization, was widely expressed in all niches, both in the extracellular matrix and in osteoblastic cells. We also noted osteonectin expression in PCa cells themselves. Thus, co-culture with PCa cells was compatible with maintaining osteoblastic and osteoclastic populations in OBCNs.

Gene expression analysis of non-cancerous cells isolated from OBCNs revealed dynamic modulation of bone-related markers in both osteoblasts and osteoclasts following co-culture with PCa cells (Figure 3B, S. Figure 6). IBSP expression was downregulated in both L- and P-OBCNs at TP1 (4 days after seeding of PCa cells) compared to cancer-free OBCNs, with the most pronounced decrease observed in P-OBCNs (9-fold lower in BM267, *p* < 0.05, Figure 3B; 4-fold lower in BM256, n.s.; S. Figure 6A). By TP2 (10 days after seeding of PCa cells) IBSP expression was undetectable across all conditions, possibly due to the absence of osteogenic factors during this phase of cell culture (S. Figure 6B, 6C). BGLAP and BMP2 expression remained rather stable across most conditions, with some donor-related variability.

For OPN and TRAP expression, we observed differential effects between L-OBCNs and P-OBCNs, with stronger downregulation of both markers in the latter. TRAP and OPN were consistently less expressed in P-OBCNs relative to OBCN controls, with ∼6.5- and ∼13-fold decreases in BM267 P-OBCNs at TP1. This suppression persisted at TP2 across donors and conditions, with a reduction ranging from ∼3.5-fold to ∼10-fold (Figure 3B, S. Figure 6A-6C). A comparable trend was observed in BM256: effects were modest at TP1 (1.2-fold, n.s., and 2.3-fold, p < 0.005; S. Figure 6A), and more pronounced at TP2 (3.5-fold, p < 0.005, and 10-fold, p < 0.05; S. Figure 6A), where both markers were significantly reduced in P-OBCNs. In contrast, co-culture with LNCaP cells (L-OBCNs) induced more modest changes in OPN and TRAP expression, which were generally not statistically significant.

Together, these results indicate that co-culture with PCa cells alters the transcriptional profile of the bone niche, with modest and variable effects on osteoblastic genes and a more consistent suppression of osteoclastic gene expression, particularly in the presence of PC3 cells.

### 2.4. Osteoclast presence in the bone niche enhances the osteoclastic gene signature of prostate cancer cells

We next asked whether PCa cell phenotype is modulated by culture in engineered niches. Specifically, we first assessed whether tumor cells exposed to OBNs or OBCNs acquired features of bone lineage cells, a process known as osteomimicry (Figure 3C, S. Figure 7A-7C). Cells cultured in 2D under identical medium conditions served as controls. Among genes associated with osteoblastic differentiation and possible markers of osteomimicry, we prioritized the analysis of BGLAP, BMP2, and SPARC (osteonectin), the latter based on its dual role in bone remodeling and PCa progression [28]. IBSP was not included in this analysis due to insufficient RNA yield from the sorted tumor cell population in these specific experiments.

In 2D cultures, baseline SPARC expression in PC3 cells was approximately 10-fold higher than in LNCaP (*p* < 0.005), suggesting a cell line-specific response to OCM. After 4 days of co-culture in engineered niches, SPARC expression was consistently upregulated (up to 9-fold, *p* < 0.05) in LNCaP cells as compared to 2D controls (Figure 3C, S. Figure 7). In PC3 cells, SPARC was also elevated, but only in BM267-based OBCNs (∼5.3-fold; *p* < 0.05). No consistent differences in SPARC expression by PCa cells cultured in OBN and OBCN were measured. In 2D cultures, BGLAP expression was either undetectable or near the detection limit (maximum observed: 4.5x10⁻⁶) across both cell lines and timepoints, precluding relevant comparison with engineered niche conditions. After co-culture in engineered niches, BGLAP was expressed at low or undetectable levels in LNCaP across donors and timepoints. In contrast, PC3 generally expressed higher levels of BGLAP across most conditions. No statistically significant differences were observed between OBN and OBCN using either cell line (Figure 3C and S. Figure 7). For 2D conditions, BMP2 expression was largely undetectable in LNCaP, while PC3 showed modest expression only at TP1 (1.5x10⁻⁴). In PCa co-cultured within niches, BMP2 expression remained below detection in all LNCaP samples across donors and timepoints. In contrast, PC3 consistently expressed BMP2 in all 3D conditions (range: 2.7x10⁻⁴ to 8.0x10⁻⁴), suggesting that OBN and OBCN support or sustain BMP2 expression specifically in this cell line (Figure 3C and S. Figure 7). Taken together, expression patterns of these markers indicate a cell line-specific response to the bone-like niche in osteoclastogenic conditions: PC3, but not LNCaP, consistently induced the osteoblast-associated markers BGLAP and BMP2, while SPARC was upregulated in both cell lines in 3D, albeit with donor- and condition-dependent variability.

Genes associated with osteoclastic differentiation and activity, namely OPN and TRAP, showed the most significant changes across conditions. In 2D cultures, expression was either undetectable or near the detection limit (maximum observed: 2.3x10⁻⁶) for both PC3 and LNCaP across all donors and timepoints. Upon co-culture in engineered niches, OPN was detected in both cell lines but with markedly higher expression in PC3, particularly under OBCN conditions, where it showed a ∼68-to >2700-fold increase in expression compared to OBNs, depending on donor and timepoint (*p* < 0.05, except BM267 TP1: *p* = 0.07). In LNCaP, OPN expression ranged from 1.8x10⁻⁶ to 2.1x10⁻³, with the highest levels observed in OBCN at TP2 in both donors. TRAP expression was similarly induced in PC3 exclusively in OBCN conditions (range: 9.0x10⁻⁵ to 7.9x10⁻⁴), with no detectable expression in OBN. In contrast, LNCaP cells expressed TRAP at low but detectable levels in both OBN and OBCN (range: 6.4x10⁻⁶ to 5.9x10⁻⁵), with a consistent 2-to 6-fold increase in OBCN compared to OBN. No statistically significant differences were observed between donors or timepoints, though niche-specific trends were consistent across replicates (Figure 3C and S. Figure 7). Together, these findings indicate a niche- and cell line-specific upregulation of osteoclast-associated genes, with PC3 cells exhibiting strong induction of OPN and TRAP exclusively in OBCN, while LNCaP showed lower but consistently enhanced expression of both markers in OBCN compared to OBN cultured in osteoclastogenic medium.

In summary, both PCa cell lines exhibited niche-induced osteomimicry, with upregulation of genes associated with osteoblastic and osteoclastic differentiation. The response was largest in PC3 cells cultured in the presence of osteoclastic cells (OBCN). Our findings indicate that the inclusion of osteoclasts within the engineered bone niche has a specific and measurable effect on PCa osteomimicry.

**Figure 3.**
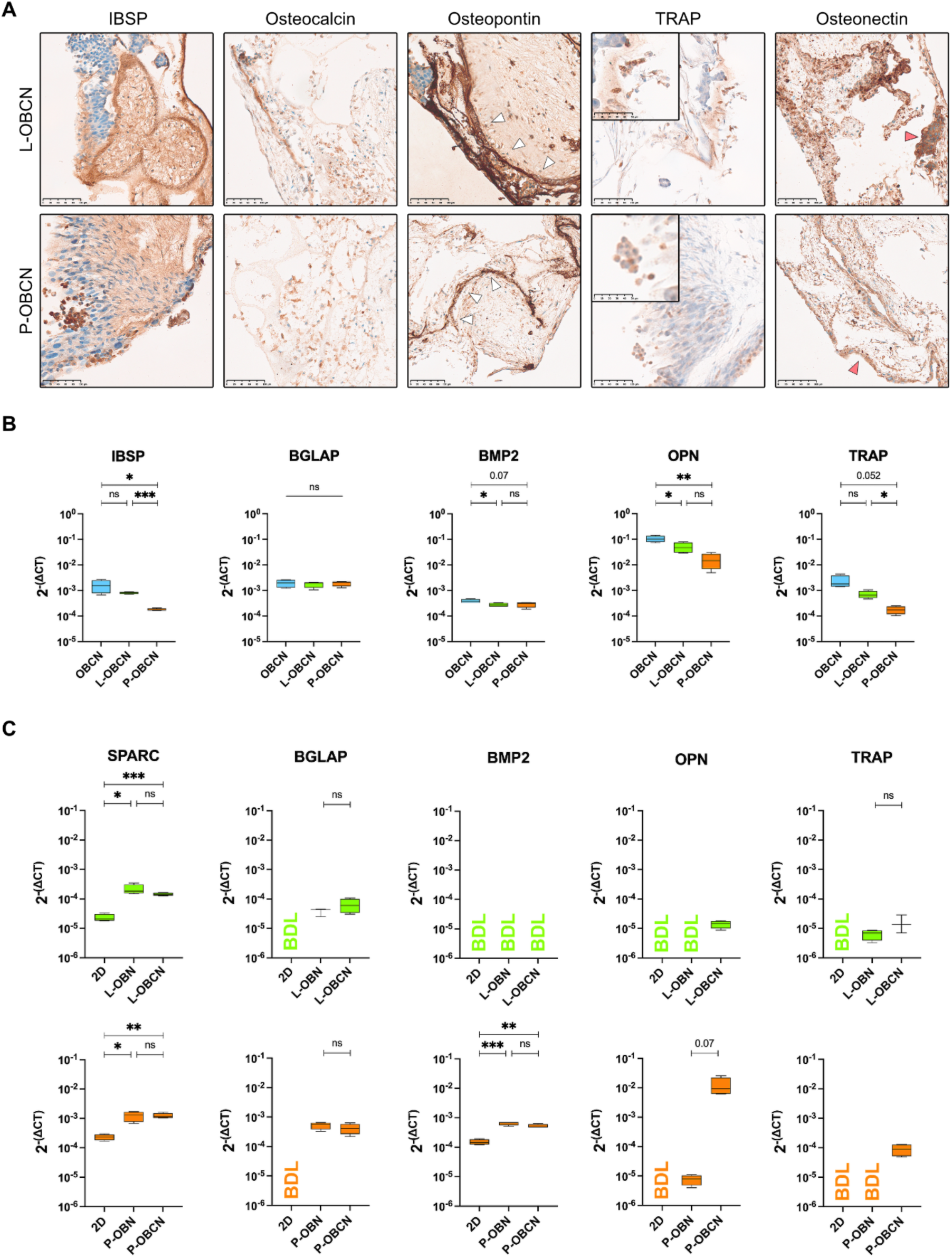
Reciprocal effects of PCa cells and engineered bone niches upon co-culture. (A) IHC imaging comparing expression of markers IBSP, osteocalcin, osteopontin, TRAP, and Osteonectin in L-OBCNs and P-OBCNs (Scale bar: 100 µm). White arrowheads in osteopontin images indicate areas adjacent to scaffold-derived hydroxyapatite. Red arrowheads show identified PCa cells expressing osteonectin. Higher-magnification images of TRAP staining are shown, highlighting the presence of multinucleated TRAP+ cells. (Scale bar: 50 µm). (B) Gene expression analysis of bone-related markers IBSP, BGLAP, BMP2, OPN, and TRAP in non-tumor niche cells (FACS-sorted). Comparisons are drawn between cells extracted from OBCNs without PCa (blue), L-OBCNs (green), and P-OBCNs (orange). Data for cancer-free OBCNs are referenced in Figure 1B. (C) qRT-PCR analysis of osteoblastic and osteoclastic markers SPARC, BGLAP, BMP2, OPN, and TRAP on PCa cells extracted from L-OBNs and L-OBCNs (green) or P-OBNs and P-OBCNs (orange). Comparisons are drawn between cells co-cultured in the niches and the same PCa line grown in 2D under the same medium conditions. Each graph represents data from a singular donor (BM267 at TP1), with up to n=4 experimental replicates. Comparisons were done via Welch t-test, with statistically significant changes being shown in the graph (✱: p < 0.05; ✱✱: p < 0.005; ✱✱✱: p < 0.0005). Conditions with undetectable gene expression have been marked with BDL (below detection level).

### 2.5. PCa patient-derived organoids successfully engraft in OBCNs and exhibit osteomimicry

We next tested whether OBCNs could support and maintain patient-derived tumor material, extending its relevance toward translational applications. We co-cultured a PCa patient-derived organoid-to-xenograft-to-organoid model (PDOXO; derived from the hormone-naïve lung metastasis resection specimen [29]) in the niche following the same protocol used for LNCaP and PC3 cells. OBCNs were then assessed for PCa colonization, growth, and bone-related marker expression by immunofluorescence staining, using formalin-fixed paraffin-embedded (FFPE) tissues of human PCa bone metastasis samples as controls.

PCa cells were identified by Cytokeratin-8 (CK8) staining at both time points and organized into clusters of varying sizes (Figure 4A). Rather than aggregating in 3D, organoid-like structures, cancer cells adhered to the niche and spread in monolayer-like clusters (S. Videos 1 and 2). The number and volume of isolated aggregates increased slightly at 10 days (TP2) compared to 4 days (TP1; Figure 4B, *p* = 6.2 x 10^-10^), demonstrating that the engineered niches can support the short-term culture of patient-derived PCa cells for up to 10 days. Notably, the same organoid line failed to grow in Matrigel under identical osteoclastogenic medium conditions, resulting in no analyzable material.

To verify that our model reproduces the inherent osteomimetic phenotype of mPCa cells, we stained the niches for osteocalcin, TRAP, and osteopontin, the markers most strongly modulated in LNCaP and PC3 co-culture. The CK8-positive organoid cells were positive for all three markers, with membrane-localized osteocalcin and TRAP expression and co-expression of osteopontin observed at both time points (Figure 4C).

We next examined whether these bone-related markers were expressed PCa bone metastasis resections. In all bone metastasis samples, CK8+ cells consistently expressed osteocalcin and TRAP. Osteopontin expression was only detected in one sample, while CK8-positive cells were adjacent to, but did not overlap with, osteopontin-positive areas in the two other samples.

These findings offer a proof-of-principle that OBCNs can host and maintain PCa patient-derived organoids *in vitro* without the need for complex media formulations. Furthermore, patient-derived PCa cells express bone-related markers within the niche, recapitulating the osteomimicry phenotype observed in both immortalized cell lines and patient samples.

**Figure 4.**
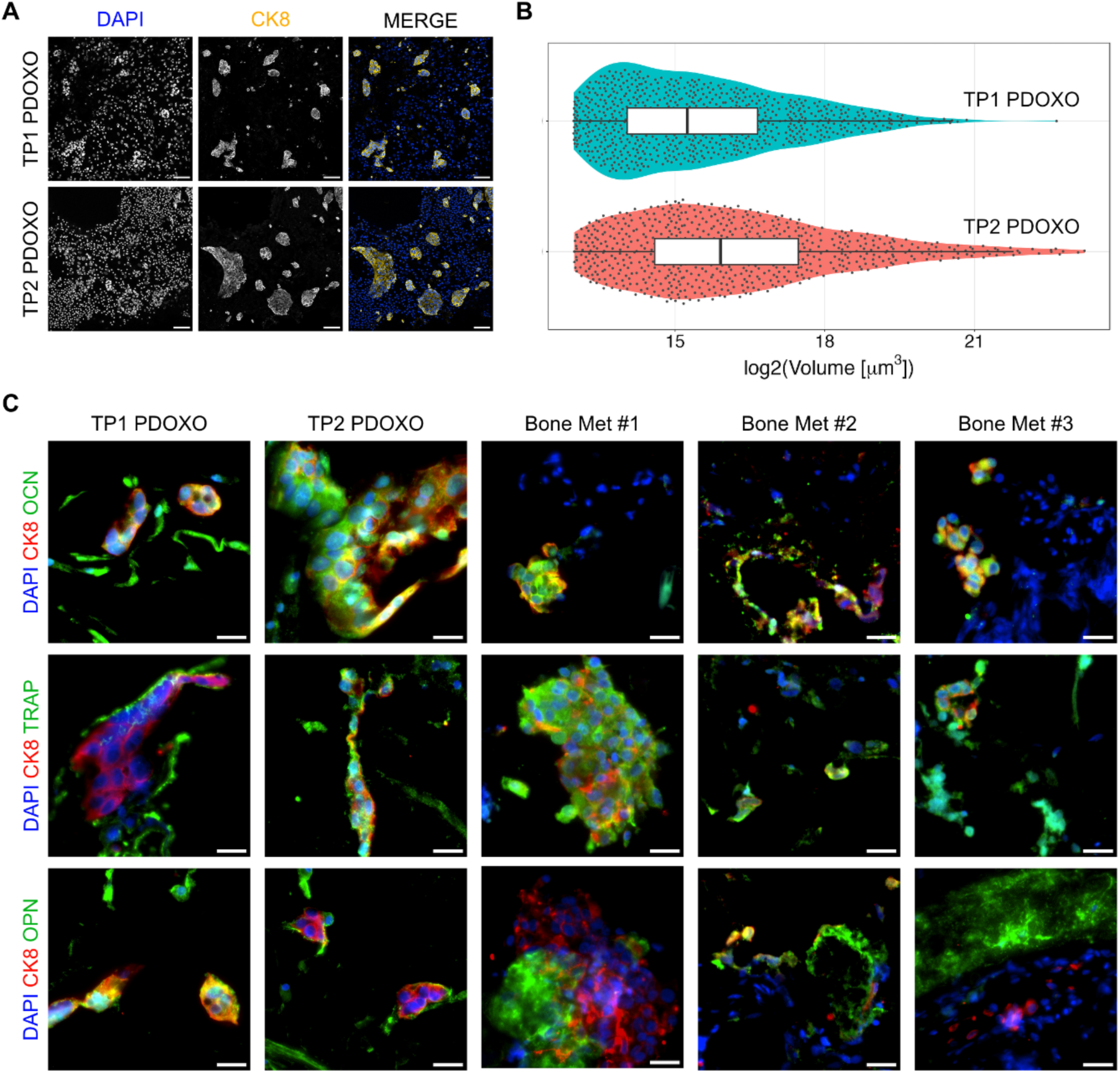
Engraftment of a patient-derived organoid line into OBCNs and maintenance of its osteomimetic behavior. (A) Immunofluorescence images showing aggregates of PCa cells from a treatment naïve lung metastasis PDX-to-PDO line, stained with DAPI (blue) and Cytokeratin-8 (yellow). Images are taken at 4 days and 10 days after the start of culture in OBCNs (Scale bar: 100 µm). (B) Distribution of CK8+ aggregate volume and their numbers for P20-11 PDOs cultured for TP1 (blue) and TP2 (red). Volumes are represented as log_2_ values. Violin plots are overlaid with the median volume value. Log median difference between distributions = 0.621. Difference in median value statistically significant per Mann-Whitney U Test (p = 6.2 x 10^-10^). (C) Immunofluorescence images showing expression of bone-related markers osteocalcin, TRAP, and osteopontin in PCa organoids cultured for 4 days and 10 days, alongside patient-derived control samples from PCa bone metastasis (Scale bar: 20 µm).

## 3. Discussion

We report the engineering of a dual-lineage (i.e., osteoblastic and osteoclastic) bone-mimicking niche supporting engraftment of PCa cell lines and patient-derived organoids. The fully human model enables the identification of reciprocal phenotypic changes in co-cultured tumor and bone cells, as well as osteoclast-regulated osteomimicry in PCa cells.

A key innovation of our approach is the integration of osteoclasts and their derivation thanks to co-culture with osteoblasts. While osteoclasts are often considered secondary to the osteoblastic predominance of metastatic lesions, they play essential roles in initiating niche remodeling, releasing matrix-derived growth factors, and modulating osteoblast activity [13–15]. Previous *in vitro* systems have either entirely excluded osteoclasts or forced their differentiation with exogenous RANKL, thereby bypassing endogenous signaling and potentially distorting niche biology. By restoring the osteoblast-osteoclast axis, our model captures physiological homeostatic processes, as a basis to investigate how osteoclast activity contributes to the metastatic bone microenvironment.

The successful differentiation of both osteoblastic and osteoclastic lineages within the OBCNs was obtained through a sequential, modular approach used in constructing the niches. Gene and protein expression of canonical osteoblastic markers (IBSP, BGLAP, BMP2, ALPL) confirmed osteoblastic maturation across donors, despite some variability. ALPL and BMP2 were maintained, while BGLAP was consistently downregulated, and IBSP showed donor-specific modulation. This pattern mirrors *in vivo* bone remodeling compartments, where osteoblast activity is preserved but terminal maturation is suppressed during active resorption. Osteoclast-derived factors such as SEMA4D and S1P have been reported to inhibit osteoblast maturation under these conditions [30], suggesting that similar mechanisms may underlie the transcriptional modulation observed in our dual-lineage niches. Osteoclastogenesis was confirmed by TRAP upregulation and the presence of multinucleated TRAP⁺ cells, consistent with Vitamin D3-driven, osteoblast-mediated RANKL induction [31,32]. OPN transcripts were also enriched in OBCNs, aligning with its role in osteoclast-mediated bone resorption [33,34].

Both LNCaP and PC3 cells modulated the bone-remodeling program of the OBCNs, with consistent downregulation of osteoclastic markers (OPN, TRAP) and selected osteogenic genes, particularly pronounced in PC3. This contrasts with *in vivo* models that report osteoclast activation in metastatic lesions [18,35], suggesting context-dependent suppression during early-stage adaptation. Conversely, co-culture in OBCNs induced a robust osteomimicry response in tumor cells, with upregulation of both osteoblastic (SPARC, BGLAP) and osteoclastic (OPN, TRAP) markers compared to 2D culture, consistent with reports linking these phenotypes to metastatic adaptation and progression [36–39]. This dual osteoblastic–osteoclastic mimicry was more marked in PC3, consistent with their bone-metastatic origin, and underscores the reciprocal, cell-line-specific interactions between PCa cells and the human bone niche.

To investigate whether osteoclasts specifically modulate the osteomimicry response, we compared PCa cells cultured in osteoblast-only niches (OBNs) versus osteoblast–osteoclast niches (OBCNs). While both niche types induced similar levels of osteoblastic markers in PCa, only OBCNs consistently drove higher expression of the osteoclastic markers OPN and TRAP. This indicates that osteoclasts enhance the osteoclastic component of the mimicry program, consistent with prior reports linking osteoclast activity to increased OPN expression in cancer cells [40].

The OBCN model also supported the engraftment and maintenance of a PCa PDOXO model initially generated from a lung metastasis. Colonization of luminal PCa cells was already evident at 4 days (TP1), when single PDOXO cells had formed large aggregates attached to the scaffold, indicating successful adhesion, cell-cell association, and potential growth. By 10 days (TP2), the median aggregate volume had further increased. The novelty of the finding is underscored by the low success rates of PCa organoid cultures [41] and the very limited number of studies achieving co-culture of patient-derived PCa cells in 3D mineralized bone-like scaffolds, exemplified by Choudhary et al. (2018), Shokoohmand et al. (2019), and Paindelli et al. (2024) [42–44]. Moreover, none of these prior studies combined patient-derived PCa cells with an endogenous osteoclastogenesis driven by osteoblasts, as implemented here.

The OBCN model thus represents a human-derived system that captures key features of the endosteal niche, reciprocal molecular modulation between tumor and bone cells, and supports patient-derived material. Moreover, its design allows for modular integration and removal of specific niche components. Considering our previously engineered environments using the same scaffold, including perivascular components and supporting hematopoietic stem and progenitor cell culture [26,27], we envision extending the approach by integrating multiple niche types into a modular, compartmentalized bone metastasis platform. This combination provides a foundation for studying targeted dissection of niche-specific mechanisms and testing of therapies that disrupt the spatial dependencies driving metastatic PCa [45–47]. The system could also support personalised drug testing in patient-specific contexts.

A key limitation of the current configuration is the short co-culture window, here limited to 10 days, dictated by the finite lifespan of osteoclasts *in vitro*. This design ensures that molecular changes reflect interactions with a niche containing intact osteoclasts, but also limits the study to early stages of tumor-bone adaptation, potentially explaining why transcriptional modulation of osteoblastic and osteoclastic markers (e.g., IBSP, TRAP, OPN) was not mirrored at the protein level. Moreover, testing a larger panel of patient-derived organoids from different metastatic sites will be necessary to determine how broadly the model applies across disease subtypes. Finally, introducing scaffolds with tunable stiffness and mineral composition will extend the model’s reach to the biomechanical heterogeneity of bone, offering new opportunities to explore how mechanical and biochemical signals converge to drive disease progression.

## 4. Conclusion

This study presents a fully human, engineered mPCa model in bone and supports co-culture with PCa cells from immortalized lines and patient-derived organoid models. The model reproduces key early features of metastatic establishment, including alterations in the bone microenvironment phenotype and the induction of dual osteoblastic-osteoclastic osteomimicry in tumor cells. Its compatibility with patient-derived material highlights potential applications in personalized drug testing and mechanistic studies of tumor-niche interactions. With further refinement this platform could help elucidate the cellular and molecular mechanisms driving metastatic PCa adaptation to bone. Finally, it could support the development of niche-targeted therapeutic strategies that bridge tissue engineering, oncology, and personalized medicine.

## 5. Methods

### 5.1. Cell culture media

Complete Medium (CM) consisted of Minimum Essential Medium Alpha (aMEM) supplemented with fetal bovine serum (FBS; same lot across all experiments), GlutaMAX, Penicillin-Streptomicin, HEPES, Sodium Pyruvate (detailed composition provided in Supplementary Information (SI) file). Unless otherwise stated, all reagents were used at working concentrations.

Proliferative Medium (PM) was prepared by supplementing CM with 5 ng/mL human fibroblast growth factor 2 (FGF2). Osteoblastic Medium (OM) was prepared by supplementing CM with 0.1 mM ascorbic acid, 10 nM dexamethasone, and 10 mM beta-glycerophosphate to promote osteogenic differentiation. Osteoclastic Medium (OCM) was composed of CM supplemented with 10 nM 1,25-dihydroxyvitamin D3 (Vitamin D3) and 25 ng/mL human macrophage colony-stimulating factor (detailed composition provided in SI file). This formulation is based on established osteoclastogenesis-inducing conditions and leverages Vitamin D3’s role in upregulating RANKL expression in osteoblasts [25].

### 5.2. Cell and tissue sources and cell expansion

Human bone marrow mesenchymal stromal cells (BM-MSCs) were isolated from bone marrow aspirates obtained during routine orthopedic surgery involving iliac crest exposure, as previously described [48]. BM-MSCs from three independent donors were used in this study: BM267, BM256, and BM293. Cells were expanded in 2D culture by seeding at a density of 5x10^3^ cells/cm^2^ in PM, with medium changes twice weekly. Cells were passaged upon reaching 80% confluence. BM-MSCs were either used fresh or cryopreserved at passage 2 (P2); for all experiments, cells were used at passage 2 or 3 (P2–P3).

Human CD14+ monocytes were obtained from peripheral blood collected through routine blood donations at the Swiss Red Cross (SRC) Blood Donation Center of Basel (University Hospital of Basel). Peripheral blood mononuclear cells (PBMCs) were isolated by density gradient centrifugation over Ficoll-Paque PLUS at 400×g for 25 minutes at room temperature without brakes. CD14⁺ cells were subsequently purified from the PBMC fraction using magnetic-activated cell sorting (MACS) with CD14 MicroBeads, LS columns, and a QuadroMACS Separator, according to the manufacturer’s protocol.

PCa cell lines LNCaP (AR+ HSPC-like) and PC3 (AR− CRPC-like, bone-derived) were cultured in standard 2D conditions by seeding at a density of 5x10^3^ cells/cm^2^ in CM. Medium was changed twice per week, and cells were passaged when they reached 80% confluence. LNCaP and PC3 cells were stably labeled with mEmerald and mCherry fluorescent reporters, respectively (Supplementary Methods (SM) for full protocol).

Patient-derived organoids (P20-11 PDOs) were established from a treatment-naïve lung metastasis specimen [29]. PDO-xenografts (PDOXs) were obtained by subcutaneous injection of dissociated PDO cells in NSG male mice in the context of another study and biobanked for further applications. PDOX tumors were then dissociated and cultured as organoids (PDOXOs) as described by Dolgos et al. (2025) [49]. Early-passage PDOXOs were maintained in prostate organoid medium (POM; detailed composition provided in [29] and SI file), enzymatically dissociated into small aggregates, and seeded directly onto pre-established osteoblastic/osteoclastic niches.

PCa bone metastasis samples (P20-28 CRCP, P22-42 under ADT, P24-08 HSPC; all derived from biopsy of vertebra adenocarcinoma) were used for immunofluorescence controls and were obtained under approval by the Ethics Committee of Northwestern and Central Switzerland (EKNZ 37/13) with informed patient consent in the context of previous studies [29,50].

### 5.3. 3D niche engineering and co-culture with PCa cells

#### 5.3.1. 3D Niche Engineering and Osteogenic Differentiation

Osteoblastic niches (OBN) and osteoblastic–osteoclastic niches (OBCN) were established under static conditions, as illustrated in Figure 1A. Briefly, a cell suspension of 5 × 10⁵ BM-MSCs in 60 µL of CM was pipetted onto custom porous hydroxyapatite scaffolds (8 mm diameter × 4 mm height), which were then placed in low-attachment 12-well plates. After 40 minutes of incubation at 37°C to allow for cell attachment, scaffolds were cultured in PM for 7 days, with a medium change on day 3. Suppliers and catalog numbers are provided in the SI file. Scaffolds were then cultured in OM for 21 days to promote osteogenic differentiation, with medium changes twice weekly.

#### 5.3.2. Seeding of Osteoclastic Precursors and Dual-Lineage Niche Formation

To generate OBCNs, OM was removed, and scaffolds were inverted to expose the scaffold face that had not been in contact with the MSC suspension. This approach ensured access to the inner pores, which are otherwise occluded by the dense MSC layer, thereby improving CD14⁺ cell infiltration. A suspension of 1x10⁶ freshly isolated CD14⁺ monocytes in 20 µL of OCM was pipetted onto the exposed surface of each scaffold and incubated for 40 minutes at 37°C. Constructs were then cultured in OCM for 14 days. OBNs underwent the same handling procedure but without CD14⁺ cell seeding.

#### 5.3.3. PCa Cell Seeding and Experimental Timeline

For PCa co-culture, transduced mEmerald-LNCaP, mCherry-PC3 cells, or organoid-derived cells (P20-11) were seeded by pipetting 1 × 10⁶ cells in 20 µL of OCM directly onto each niche after removing the medium. Following a 40-minute incubation at 37°C, scaffolds were cultured in OCM until analysis, which was performed at day 7 (timepoint 1, TP1) and day 14 (timepoint 2, TP2) after CD14⁺ addition. The co-culture duration with PCa cells was limited to 10 days after tumor cell seeding. We chose this timeline based on the *in vitro* lifespan of human monocyte-derived osteoclasts, which typically maintain peak activity for approximately 2 weeks before undergoing apoptosis or loss of resorptive function [51,52].

### 5.4. Cell isolation from 3D constructs

Niche constructs were retrieved from culture, rinsed once in PBS, and quartered using a sterile scalpel. Scaffold fragments were transferred into 1.5 mL tubes and enzymatically digested in two sequential steps: first with 0.15% Type II Collagenase at 37°C for 45 minutes with shaking at 400 rpm, followed by 0.25% Trypsin-EDTA for 15 minutes under the same conditions. This combination enabled efficient matrix degradation and single-cell release.

The resulting cell suspension was diluted 1:4 in FACS buffer (PBS supplemented with 0.5% FBS and 2 mM EDTA), centrifuged at 400 × g for 5 minutes, and resuspended in fresh buffer. To ensure a single-cell suspension, samples were filtered twice through 70 µm cell strainers. Cell preparations were used for flow cytometry and other downstream applications. Reagent sources are listed in the SI File.

### 5.5. Sorting of mEmerald-LNCaP and mCherry-PC3 cells from bone niche constructs

Cell suspensions obtained from digested scaffolds at the endpoint of culture were subjected to fluorescence-activated cell sorting (FACS) to isolate mEmerald⁺ LNCaP and mCherry⁺ PC3 cells from the total niche-derived population (Figure 2A). Gating strategies included forward and side scatter area (FSC-A vs. SSC-A), and singlet discrimination using FSC-A vs. FSC-H and SSC-A vs. SSC-W with fluorescence-based selection of the respective reporter-positive tumor populations. Control constructs without fluorescently tagged tumor cells were processed in parallel using identical gating and sorting parameters to control for potential sorting-induced effects on downstream analyzes.

### 5.6. RNA extraction, cDNA synthesis, and qRT-PCR analysis

Total RNA from sorted cell populations was extracted using either the RNeasy Mini Kit or the RNeasy Micro Kit, depending on cell yield, according to the manufacturer’s instructions. RNA concentration and purity were assessed using a NanoDrop One spectrophotometer. Complementary DNA (cDNA) was synthesized via reverse transcription using SuperScript III Reverse Transcriptase.

Quantitative real-time PCR (qRT-PCR) was performed using a ViiA 7 Real-Time PCR System and TaqMan gene expression assays targeting integrin-binding sialoprotein (*IBSP*), osteocalcin (*BGLAP*), bone morphogenic protein 2 (*BMP2*), alkaline phosphatase (*ALPL*), osteopontin (*OPN*), tartrate-resistant acid phosphatase (*TRAP*), and osteonectin (*SPARC*). *GAPDH* was used as the housekeeping gene. Relative gene expression levels were calculated using the 2^−ΔCT^ method. Suppliers and catalog numbers are provided in the SI file.

### 5.7. Whole-mount immunofluorescence staining and confocal microscopy

Scaffolds were removed from culture and fixed overnight at 4°C in 4% formaldehyde. Samples were then permeabilized and blocked for 6 hours at room temperature in PBS containing 0.4% Triton X-100, 2% bovine serum albumin (BSA), and 5% goat serum. Fixed constructs were then incubated at 4°C for 70 hours with primary antibodies against cytokeratin-8 (CK8) and tartrate-resistant acid phosphatase (TRAP), 1:200 in blocking solution. Samples were washed in PBS and incubated overnight at 4°C with secondary antibodies, Goat Anti-Mouse 546 and Goat Anti-Rabbit 647 each at 1:200 in PBS with 0.4% Triton X-100 and 2% BSA. DAPI was used as nuclear counterstain. Suppliers and catalog numbers are provided in the SI file. Samples were mounted on 35 mm imaging dishes and imaged using a Nikon Ti2-E inverted microscope equipped with an AxR point-scanning confocal unit and controlled via Nikon NIS software.

### 5.8. Histological processing, immunohistochemistry, and immunofluorescence staining

Following culture, the engineered niches were rinsed in PBS and fixed overnight in 4% formaldehyde. Samples were embedded in HistoGel and decalcified by incubation in 15% EDTA at 37°C for two weeks. Decalcified samples were embedded in paraffin and sectioned for histological analysis. Engineered tissue sections were deparaffinized, rehydrated, and stained with hematoxylin and eosin (H&E) using a standardized protocol on an automated stainer (Epredia Gemini AS). We used standard indirect immunoperoxidase procedures for IHC staining of bone sialoprotein (BSP), osteocalcin (OCN), osteopontin (OPN), osteonectin (ON), Tartrate Resistant Acid Phosphatase (TRAP), and E-Cadherin (E-Cad) on the Ventana BenchMark ULTRA automated immunostainer. Whole-slide images were acquired using a slide scanner (40x objective, NA 0.75).

For immunofluorescence, after deparaffinization, antigen retrieval was performed by incubating slides in pH 6 citrate buffer at 96°C for 15 minutes. Sections were then permeabilized in PBS containing 0.4% Triton X-100 (PBS-T) and blocked in PBS-T with 10% goat serum. Primary antibodies for cytokeratin-8 (CK8), OCN, OPN, and TRAP were incubated overnight at 1:200 in PBS-T with 1% goat serum. Sections were then incubated with secondary antibodies Goat Anti-Rabbit 488, and Goat Anti-Mouse 647, diluted 1:200 in PBS with 1% goat serum. DAPI was added for nuclear counterstaining. Images were acquired using a Nikon Ti2 widefield fluorescence microscope equipped with a Nikon DS-Ri2 camera and controlled via NIS Elements AR software. Suppliers and catalog numbers are provided in the Supplementary Information file.

Qualitative histological observations, including the morphological identification of specific cell types, was consistently supported by an expert pathologist.

### 5.9. Quantification of PCa organoid aggregates

At both timepoints, niches co-cultured with the PCa organoid line P20-11 were fixed and stained by whole-mount immunofluorescence for cytokeratin-8 (CK8), then counterstained with DAPI. Confocal z-stacks were acquired at low magnification, and we performed volumetric image analysis to quantify the number and size of CK8⁺ tumor aggregates. The data represent three independent OBCN co-cultures. Full image acquisition and analysis parameters are detailed in the Supplementary Methods.

### 5.10. Statistics

Statistical analysis of qRT-PCR data was performed using GraphPad Prism (version 10.4.2). Group comparisons were conducted using Welch’s *t*-test to account for unequal variances. All tests were two-tailed, and *p*-value < 0.05 were considered statistically significant. Tumor volumes measured at 4 (TP1) and 10 days (TP2) were log₂-transformed, and comparisons between the two time points were performed using the non-parametric Mann-Whitney U test.

## 7. Acknowledgements

We acknowledge the invaluable support of the Flow Cytometry, Microscopy, and Histology facilities at the Department of Biomedicine, the IHC facility of the Institute of Pathology (Petra Hirschmann) at the Pathology Department of the University Hospital of Basel, and the Nanoimaging facilities at the Biozentrum, University of Basel, for their expert assistance and technical contributions. We thank Dr. Romuald Parmentier for performing bioinformatic analysis on published datasets, which helped guide our analysis and discussion. We thank Dr. Igor Cervenk for analysing organoid aggregate data and related statistical analysis shown in Figure 4C. We wholeheartedly thank Associate Professor Nathalie Bock for her valuable review of the manuscript. Figures 1A and 2A were created with BioRender.com.

## 9. Data Availability statement

The data supporting the findings of this study are available within the article and its Supplementary Information files or from the corresponding author upon reasonable request.

## References

1. Cancer Today, (n.d.). https://gco.iarc.who.int/today/ (accessed July 22, 2025).

[2] R. Raychaudhuri, D.W. Lin, R.B. Montgomery, Prostate Cancer: A Review, JAMA 333 (2025) 1433–1446. 10.1001/jama.2025.0228.

[3] S. Casimiro, T.A. Guise, J. Chirgwin, The critical role of the bone microenvironment in cancer metastases, Mol. Cell. Endocrinol. 310 (2009) 71–81. 10.1016/j.mce.2009.07.004.

[4] J.-J. Body, S. Casimiro, L. Costa, Targeting bone metastases in prostate cancer: improving clinical outcome, Nat. Rev. Urol. 12 (2015) 340–356. 10.1038/nrurol.2015.90.

[5] R.B. Berish, A.N. Ali, P.G. Telmer, J.A. Ronald, H.S. Leong, Translational models of prostate cancer bone metastasis, Nat. Rev. Urol. 15 (2018) 403–421. 10.1038/s41585-018-0020-2.

[6] D.G. Hackam, D.A. Redelmeier, Translation of Research Evidence From Animals to Humans, JAMA 296 (2006) 1727. 10.1001/jama.296.14.1731.

[7] F. Salamanna, D. Contartese, M. Maglio, M. Fini, A systematic review on in vitro 3D bone metastases models: A new horizon to recapitulate the native clinical scenario?, Oncotarget 7 (2016) 44803–44820. 10.18632/oncotarget.8394.

[8] N. Cristini, M. Tavakoli, M. Sanati, S.A. Yavari, Exploring bone-tumor interactions through 3D *in vitro* models: Implications for primary and metastatic cancers, J. Bone Oncol. 53 (2025) 100698. 10.1016/j.jbo.2025.100698.

[9] C. Sánchez-de-Diego, R.C. Yada, N. Sethakorn, P.G. Geiger, A.B. Ding, E. Heninger, F. Ahmed, M. Virumbrales-Muñoz, N. Lupsa, E. Bartels, K. Stewart, S.M. Ponik, M.N. Sharifi, J.M. Lang, D.J. Beebe, S.C. Kerr, Engineering the bone metastatic prostate cancer niche through a microphysiological system to report patient-specific treatment response, Commun. Biol. 8 (2025) 961. 10.1038/s42003-025-08384-2.

[10] N. Bock, A. Shokoohmand, T. Kryza, J. Röhl, J. Meijer, P.A. Tran, C.C. Nelson, J.A. Clements, D.W. Hutmacher, Engineering osteoblastic metastases to delineate the adaptive response of androgen-deprived prostate cancer in the bone metastatic microenvironment, Bone Res. 7 (2019) 13. 10.1038/s41413-019-0049-8.

[11] A. Bessot, F. Medeiros Savi, J. Gunter, J. Mendhi, S. Amini, D. Waugh, J. McGovern, D.W. Hutmacher, N. Bock, Humanized In Vivo Bone Tissue Engineering: In Vitro Preculture Conditions Control the Structural, Cellular, and Matrix Composition of Humanized Bone Organs, Adv. Healthc. Mater. 14 (2025) 2401939. 10.1002/adhm.202401939.

[12] K.S. Koeneman, F. Yeung, L.W.K. Chung, Osteomimetic properties of prostate cancer cells: A hypothesis supporting the predilection of prostate cancer metastasis and growth in the bone environment, The Prostate 39 (1999) 246–261. 10.1002/(SICI)1097-0045(19990601)39:4%253C246::AID-PROS5%253E3.0.CO;2-U.

[13] S.-Y. Sung, C.-L. Hsieh, A. Law, H.E. Zhau, S. Pathak, A.S. Multani, S. Lim, I.M. Coleman, L.-C. Wu, W.D. Figg, W.L. Dahut, P. Nelson, J.K. Lee, M.B. Amin, R. Lyles, P.A.J. Johnstone, F.F. Marshall, L.W.K. Chung, Coevolution of Prostate Cancer and Bone Stroma in Three-Dimensional Coculture: Implications for Cancer Growth and Metastasis, Cancer Res. 68 (2008) 9996–10003. 10.1158/0008-5472.CAN-08-2492.

[14] S. Sieh, A.A. Lubik, J.A. Clements, C.C. Nelson, D.W. Hutmacher, Interactions between human osteoblasts and prostate cancer cells in a novel 3D in vitro model, Organogenesis 6 (2010) 181–188. 10.4161/org.6.3.12041.

[15] N. Bock, T. Kryza, A. Shokoohmand, J. Röhl, A. Ravichandran, M.-L. Wille, C.C. Nelson, D.W. Hutmacher, J.A. Clements, In vitro engineering of a bone metastases model allows for study of the effects of antiandrogen therapies in advanced prostate cancer, Sci. Adv. 7 (2021) eabg2564. 10.1126/sciadv.abg2564.

[16] M. Wang, F. Xia, Y. Wei, X. Wei, Molecular mechanisms and clinical management of cancer bone metastasis, Bone Res. 8 (2020) 30. 10.1038/s41413-020-00105-1.

[17] H. Yonou, A. Ochiai, M. Goya, N. Kanomata, S. Hokama, M. Morozumi, K. Sugaya, T. Hatano, Y. Ogawa, Intraosseous growth of human prostate cancer in implanted adult human bone: Relationship of prostate cancer cells to osteoclasts in osteoblastic metastatic lesions, The Prostate 58 (2004) 406–413. 10.1002/pros.10349.

[18] I. Roato, P. D’Amelio, E. Gorassini, A. Grimaldi, L. Bonello, C. Fiori, L. Delsedime, A. Tizzani, A.D. Libero, G. Isaia, R. Ferracini, Osteoclasts Are Active in Bone Forming Metastases of Prostate Cancer Patients, PLOS ONE 3 (2008) e3627. 10.1371/journal.pone.0003627.

[19] X. Chen, X. Zhi, J. Wang, J. Su, RANKL signaling in bone marrow mesenchymal stem cells negatively regulates osteoblastic bone formation, Bone Res. 6 (2018) 34. 10.1038/s41413-018-0035-6.

[20] W. Cai, Y. Huo, Y. Liu, Y. Su, H. Guo, L. Wang, B. Li, T. Liang, Biomechanics in bone regeneration and mechanobiology in osteoblasts: Fundamental concepts and recent progress, EngMedicine 2 (2025) 100057. 10.1016/j.engmed.2025.100057.

[21] K. Urano, Y. Tanaka, T. Tominari, M. Takatoya, D. Arai, S. Miyata, C. Matsumoto, C. Miyaura, Y. Numabe, Y. Itoh, M. Hirata, M. Inada, The stiffness and collagen control differentiation of osteoclasts with an altered expression of c-Src in podosome, Biochem. Biophys. Res. Commun. 704 (2024) 149636. 10.1016/j.bbrc.2024.149636.

[22] A. Matsugaki, T. Harada, Y. Kimura, A. Sekita, T. Nakano, Dynamic Collision Behavior Between Osteoblasts and Tumor Cells Regulates the Disordered Arrangement of Collagen Fiber/Apatite Crystals in Metastasized Bone, Int. J. Mol. Sci. 19 (2018) 3474. 10.3390/ijms19113474.

[23] F. Salamanna, V. Borsari, S. Brogini, G. Giavaresi, A. Parrilli, S. Cepollaro, M. Cadossi, L. Martini, A. Mazzotti, M. Fini, An in vitro 3D bone metastasis model by using a human bone tissue culture and human sex-related cancer cells, Oncotarget 7 (2016) 76966–76983. 10.18632/oncotarget.12763.

[24] S. Marino, R.T. Bishop, G. Carrasco, J.G. Logan, B. Li, A.I. Idris, Pharmacological Inhibition of NFκB Reduces Prostate Cancer Related Osteoclastogenesis In Vitro and Osteolysis Ex Vivo, Calcif. Tissue Int. 105 (2019) 193–204. 10.1007/s00223-019-00538-9.

[25] A. Papadimitropoulos, A. Scherberich, S. Güven, N. Theilgaard, H.J.A. Crooijmans, F. Santini, K. Scheffler, A. Zallone, I. Martin, A 3D in vitro bone organ model using human progenitor cells, Eur. Cell. Mater. 21 (2011) 445–458; discussion 458. 10.22203/ecm.v021a33.

[26] P.E. Bourgine, T. Klein, A.M. Paczulla, T. Shimizu, L. Kunz, K.D. Kokkaliaris, D.L. Coutu, C. Lengerke, R. Skoda, T. Schroeder, I. Martin, In vitro biomimetic engineering of a human hematopoietic niche with functional properties, Proc. Natl. Acad. Sci. U. S. A. 115 (2018) E5688–E5695. 10.1073/pnas.1805440115.

[27] G. Born, M. Nikolova, A. Scherberich, B. Treutlein, A. García-García, I. Martin, Engineering of fully humanized and vascularized 3D bone marrow niches sustaining undifferentiated human cord blood hematopoietic stem and progenitor cells, J. Tissue Eng. 12 (2021) 20417314211044855. 10.1177/20417314211044855.

[28] N. Ribeiro, S.R. Sousa, R.A. Brekken, F.J. Monteiro, Role of SPARC in Bone Remodeling and Cancer-Related Bone Metastasis, J. Cell. Biochem. 115 (2014) 17–26. 10.1002/jcb.24649.

[29] R. Servant, M. Garioni, T. Vlajnic, M. Blind, H. Pueschel, D.C. Müller, T. Zellweger, A.J. Templeton, A. Garofoli, S. Maletti, S. Piscuoglio, M.A. Rubin, H. Seifert, C.A. Rentsch, L. Bubendorf, C. Le Magnen, Prostate cancer patient-derived organoids: detailed outcome from a prospective cohort of 81 clinical specimens, J. Pathol. 254 (2021) 543–555. 10.1002/path.5698.

[30] J.-M. Kim, C. Lin, Z. Stavre, M.B. Greenblatt, J.-H. Shim, Osteoblast-Osteoclast Communication and Bone Homeostasis, Cells 9 (2020) 2073. 10.3390/cells9092073.

[31] S.-K. Lee, J. Kalinowski, S. Jastrzebski, J.A. Lorenzo, 1,25 (OH)2 Vitamin D3-Stimulated Osteoclast Formation in Spleen-Osteoblast Cocultures Is Mediated in Part by Enhanced IL-1α and Receptor Activator of NF-κB Ligand Production in Osteoblasts1, J. Immunol. 169 (2002) 2374–2380. 10.4049/jimmunol.169.5.2374.

[32] S. Kitazawa, K. Kajimoto, T. Kondo, R. Kitazawa, Vitamin D3 supports osteoclastogenesis via functional vitamin D response element of human RANKL gene promoter, J. Cell. Biochem. 89 (2003) 771–777. 10.1002/jcb.10567.

[33] M.A. Chellaiah, N. Kizer, R. Biswas, U. Alvarez, J. Strauss-Schoenberger, L. Rifas, S.R. Rittling, D.T. Denhardt, K.A. Hruska, Osteopontin Deficiency Produces Osteoclast Dysfunction Due to Reduced CD44 Surface Expression, Mol. Biol. Cell 14 (2003) 173–189. 10.1091/mbc.e02-06-0354.

[34] J. Luukkonen, M. Hilli, M. Nakamura, I. Ritamo, L. Valmu, K. Kauppinen, J. Tuukkanen, P. Lehenkari, Osteoclasts secrete osteopontin into resorption lacunae during bone resorption, Histochem. Cell Biol. 151 (2019) 475–487. 10.1007/s00418-019-01770-y.

[35] J.H. Tae, I.H. Chang, Animal models of bone metastatic prostate cancer, Investig. Clin. Urol. 64 (2023) 219–228. 10.4111/icu.20230026.

[36] N. Rucci, A. Teti, Osteomimicry: how tumor cells try to deceive the bone, Front. Biosci.-Sch. 2 (2010) 907–915. 10.2741/S110.

[37] X. Pang, R. Xie, Z. Zhang, Q. Liu, S. Wu, Y. Cui, Identification of SPP1 as an Extracellular Matrix Signature for Metastatic Castration-Resistant Prostate Cancer, Front. Oncol. 9 (2019). 10.3389/fonc.2019.00924.

[38] L.M. Adams, M.J. Warburton, A.R. Hayman, Human breast cancer cell lines and tissues express tartrate-resistant acid phosphatase (TRAP), Cell Biol. Int. 31 (2007) 191–195. 10.1016/j.cellbi.2006.09.022.

[39] S. Zenger, W. He, B. Ek-Rylander, D. Vassiliou, R. Wedin, H. Bauer, G. Andersson, Differential expression of tartrate-resistant acid phosphatase isoforms 5a and 5b by tumor and stromal cells in human metastatic bone disease, Clin. Exp. Metastasis 28 (2011) 65–73. 10.1007/s10585-010-9358-4.

[40] X. Pang, K. Gong, X. Zhang, S. Wu, Y. Cui, B.-Z. Qian, Osteopontin as a multifaceted driver of bone metastasis and drug resistance, Pharmacol. Res. 144 (2019) 235–244. 10.1016/j.phrs.2019.04.030.

[41] A. Buskin, E. Scott, R. Nelson, L. Gaughan, C.N. Robson, R. Heer, A.C. Hepburn, Engineering prostate cancer in vitro: what does it take?, Oncogene 42 (2023) 2417–2427. 10.1038/s41388-023-02776-6.

[42] S. Choudhary, P. Ramasundaram, E. Dziopa, C. Mannion, Y. Kissin, L. Tricoli, C. Albanese, W. Lee, J. Zilberberg, Human ex vivo 3D bone model recapitulates osteocyte response to metastatic prostate cancer, Sci. Rep. 8 (2018) 17975. 10.1038/s41598-018-36424-x.

[43] A. Shokoohmand, J. Ren, J. Baldwin, A. Atack, A. Shafiee, C. Theodoropoulos, M.-L. Wille, P.A. Tran, L.J. Bray, D. Smith, N. Chetty, P.M. Pollock, D.W. Hutmacher, J.A. Clements, E.D. Williams, N. Bock, Microenvironment engineering of osteoblastic bone metastases reveals osteomimicry of patient-derived prostate cancer xenografts, Biomaterials 220 (2019) 119402. 10.1016/j.biomaterials.2019.119402.

[44] C. Paindelli, V. Parietti, S. Barrios, P. Shepherd, T. Pan, W.-L. Wang, R.L. Satcher, C.J. Logothetis, N. Navone, M.T. Campbell, A.G. Mikos, E. Dondossola, Bone mimetic environments support engineering, propagation, and analysis of therapeutic response of patient-derived cells, *ex vivo* and *in vivo*, Acta Biomater. 178 (2024) 83–92. 10.1016/j.actbio.2024.02.025.

[45] S.H. Park, E.T. Keller, Y. Shiozawa, Bone Marrow Microenvironment as a Regulator and Therapeutic Target for Prostate Cancer Bone Metastasis, Calcif. Tissue Int. 102 (2018) 152–162. 10.1007/s00223-017-0350-8.

[46] M.E. Sowder, R.W. Johnson, Bone as a Preferential Site for Metastasis, JBMR Plus 3 (2019) e10126. 10.1002/jbm4.10126.

47. E.C. Kabak, S.L. Foo, M. Rafaeva, I. Martin, M. Bentires-Alj, Microenvironmental Regulation of Dormancy in Breast Cancer Metastasis: “An Ally that Changes Allegiances,” in: T. Sørlie, R.B. Clarke (Eds.), Guide Breast Cancer Res. Cell. Heterog. Mol. Mech. Ther., Springer Nature Switzerland, Cham, 2025: pp. 373–395. 10.1007/978-3-031-70875-6_18.

[48] N. Sadr, B.E. Pippenger, A. Scherberich, D. Wendt, S. Mantero, I. Martin, A. Papadimitropoulos, Enhancing the biological performance of synthetic polymeric materials by decoration with engineered, decellularized extracellular matrix, Biomaterials 33 (2012) 5085–5093. 10.1016/j.biomaterials.2012.03.082.

[49] R. Dolgos, R. Parmentier, J. Wang, R. Servant, A.J. Templeton, T. Zellweger, A.D. Lamb, K.D. Mertz, S. Subotic, T. Vlajnic, H. Seifert, A. Mortezavi, C.A. Rentsch, L. Bubendorf, C. Le Magnen, Single-cell analysis uncovers preserved prostate cancer lineages and universally altered pathways in Matrigel-free patient-derived organoids, Cell Rep. 44 (2025) 116352. 10.1016/j.celrep.2025.116352.

50. R. Dolgos, R. Parmentier, J. Wang, R. Servant, A.J. Templeton, T. Zellweger, A.D. Lamb, K.D. Mertz, S. Subotic, T. Vlajnic, H. Seifert, A. Mortezavi, C.A. Rentsch, L. Bubendorf, C.L. Magnen, ECM-free patient-derived organoids preserve diverse prostate cancer lineages and uncover in vitro-enriched cell types, (2024) 2024.10.16.618617. 10.1101/2024.10.16.618617.

[51] K. Fujisaki, N. Tanabe, N. Suzuki, T. Kawato, O. Takeichi, O. Tsuzukibashi, M. Makimura, K. Ito, M. Maeno, Receptor activator of NF-κB ligand induces the expression of carbonic anhydrase II, cathepsin K, and matrix metalloproteinase-9 in osteoclast precursor RAW264.7 cells, Life Sci. 80 (2007) 1311–1318. 10.1016/j.lfs.2006.12.037.

[52] E. Rossi, E. Mracsko, A. Papadimitropoulos, N. Allafi, D. Reinhardt, A. Mehrkens, I. Martin, I. Knuesel, A. Scherberich, An In Vitro Bone Model to Investigate the Role of Triggering Receptor Expressed on Myeloid Cells-2 in Bone Homeostasis, Tissue Eng. Part C Methods 24 (2018) 391–398. 10.1089/ten.tec.2018.0061.

