## Supplementary Material for "Dual-Lineage Human 3D Bone Niche Model Reveals Osteoclast-Driven Osteomimicry in Prostate Cancer"

### 1. Supplementary Methods

#### 1.1. Lentiviral labeling of PCa cell lines

LNCaP and PC3 cells were stably labeled with mEmerald and mCherry fluorescent reporters, respectively, using third-generation lentiviral vectors based on the pLVX backbone and carrying a neomycin resistance cassette. Lentiviral particles were produced in Lenti-X 293T cells using standard packaging plasmids and Lipofectamine 2000. Cells were transduced at a multiplicity of infection (MOI) of 2 in the presence of polybrene, and selected with G418 for 10 days. Stable expression was confirmed by fluorescence microscopy before co-culture. Suppliers and catalog numbers are provided in the Supplementary Information file.

#### 1.2. FACS gating and instrument parameters

Sorting was performed using either a CytoFLEX SRT (Beckman Coulter) or FACSAria III (BD Biosciences), depending on instrument availability. mEmerald$^+$ LNCaP cells were detected using the green channel (B525/40 on the CytoFLEX SRT; 488–530/30 on the FACSAria III), and mCherry$^+$ PC3 cells were detected using the red channel (Y610/20 on the CytoFLEX SRT; 561–616/23 on the FACSAria III). Initial gating was based on FSC-A vs. SSC-A, followed by singlet discrimination using FSC-A vs. FSC-H and SSC-A vs. SSC-W. Fluorescence compensation and gating thresholds were established using single-color controls and non-fluorescent negative samples processed identically to the experimental groups.

#### 1.3. Whole-mount immunofluorescence confocal acquisition parameters

Samples were mounted in 35 mm imaging dishes (Ibidi) and imaged using a Nikon Ti2-E inverted microscope equipped with an AxR point-scanning confocal unit and NIS-Elements software. Fluorescence excitation was performed using 405, 488, 561, and 640 nm lasers, with emission detection windows at 429–474, 499–525, 571–605, and 662–737 nm, respectively. Images were acquired in 12-bit format using resonant scanning mode with 16× line averaging, 2048 × 2048 pixel resolution, and a 1 Airy unit pinhole (16 μm at 640 nm). Z-stacks were collected across 50–600 μm depths using AutoSignal.ai to optimize laser power and detector gain. Step intervals were 11.6 μm (4×), 2.1 μm (10×), 1.0 μm (20×), and 0.3 μm (40×) objectives.

#### 1.4. Image acquisition and volumetric quantification of organoid aggregates

Confocal z-stacks were acquired on a Nikon Ti2-E microscope equipped with an AxR confocal scanning unit, using a 4× objective. Z-stacks consisted of 27 to 52 optical sections with a z-step of 11.6 μm.

Image stacks were converted to volumetric .ims format using Imaris File Converter 10.1.0 and analyzed in Imaris 10.1.0 (Bitplane) using the Measurement Pro module. Surfaces were generated from the CK8 channel via absolute intensity thresholding, using the default surface reconstruction algorithm. Objects with a volume smaller than $8 \times 10^3$ μm$^3$ were excluded to eliminate background signal and debris. For each construct, the number and volume of $CK8^+$ tumor aggregates were quantified and exported for downstream statistical analysis.

#### 1.5. Scanning electron microscopy (SEM)

At each timepoint, bone niche constructs were fixed overnight in 2% glutaraldehyde prepared in 0.1 M sodium cacodylate buffer (pH 7.4). Samples were dehydrated through a graded ethanol series (30%, 50%, 70%, 90%, and 100%) with 15-minute incubations at each step. Dehydrated constructs were processed using a critical point dryer (Leica EM CPD300) and mounted on aluminum stubs using carbon adhesive. Samples were sputter-coated with a 20 nm layer of gold using a Leica EM ACE600 coater. SEM images were acquired using a ZEISS Gemini 2 field-emission scanning electron microscope operating at 5 kV accelerating voltage and 200 pA probe current. Catalog numbers and reagent details are listed in the SI File.

### 2. Supplementary Figures

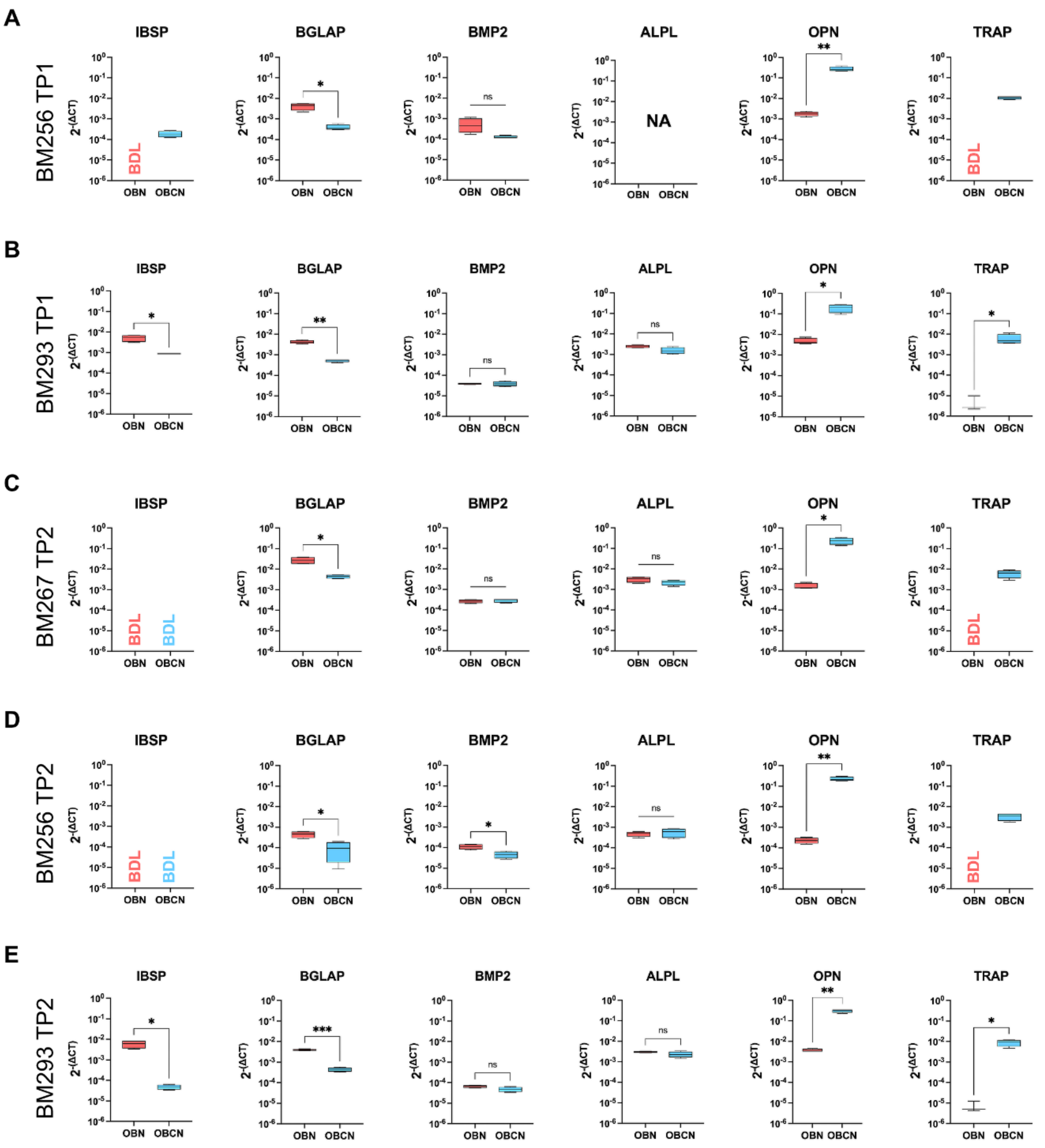

**Supplementary Figure 1 - Osteoblastic and osteoclastic marker expression compared between OBNs (red) and OBCNs (blue) by qRT-PCR analysis.** The analysis was performed for markers IBSP, BGLAP, BMP2, ALPL, OPN, and TRAP.

Analysis was done for the following donors and conditions: (A) donor BM256 at TP1, (B) BM293 at TP1, (C) BM267 at TP2, (D) BM256 at TP2, (E) BM293 at TP2. Expression level was normalized to GAPDH and calculated using the $2^{-\Delta CT}$ method. Each graph represents data from a singular donor, at the first analysis timepoint, with up to n=4 experimental replicates. Comparisons were done via Welch t-test, with statistically significant changes being shown in the graph (✱: $p < 0.05$; ✱✱: $p < 0.005$; ✱✱✱: $p < 0.0005$). Undetectable levels of gene expression were marked as BDL (below detection level), while ALPL for BM256 at TP1 could not be assessed and was marked as NA (not available).

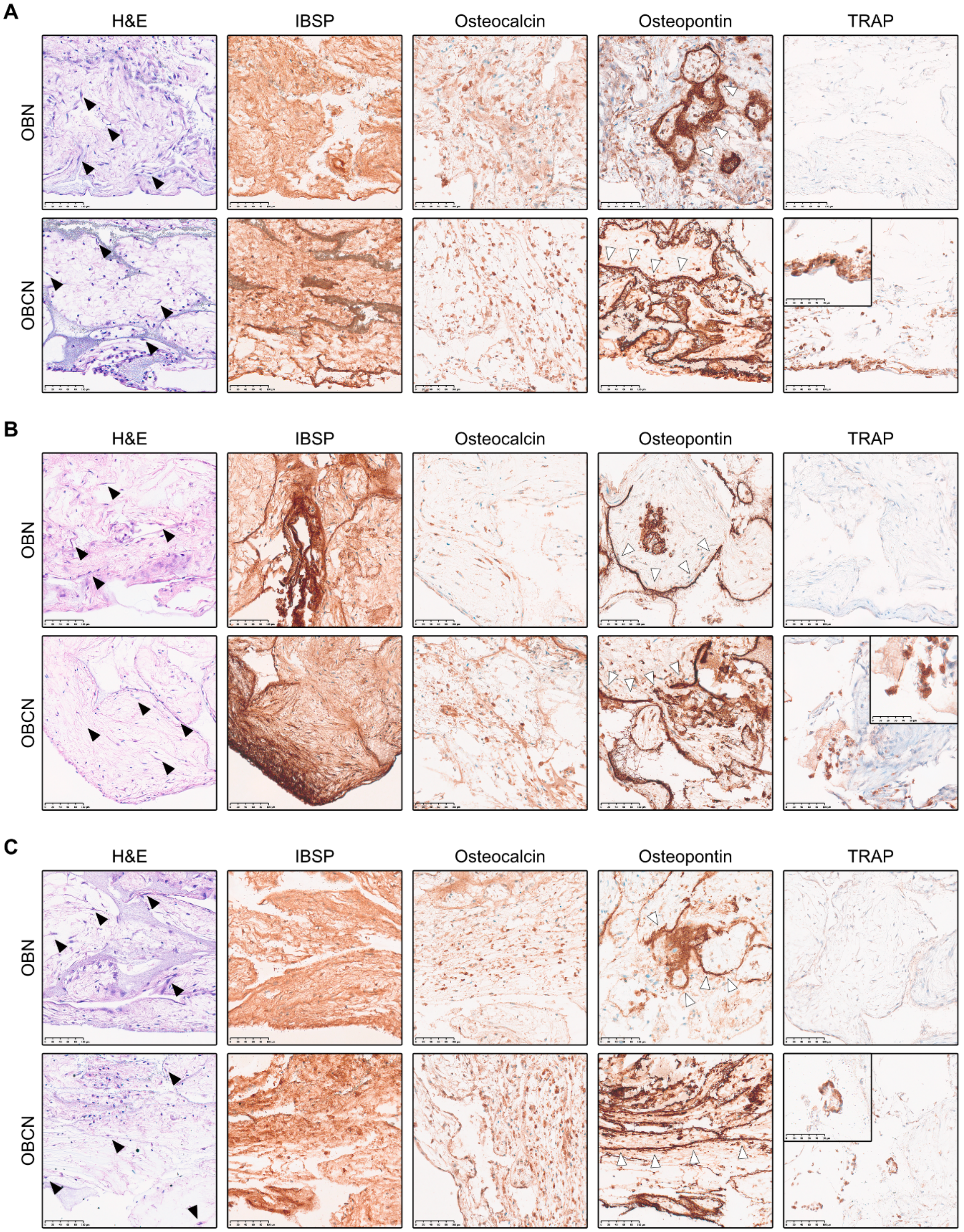

**Supplementary Figure 2 - Immunohistochemistry (IHC) imaging of osteoblastic and osteoclastic markers in OBNs and OBCNs.** Niches were imaged via H&E staining and IHC for IBSP, osteocalcin, osteopontin, and TRAP (Scale bars: 100 μm). Imaging was done for donors (A) BM256 at TP1, (B) BM267 at TP2, and (C) BM256 at TP2. Black arrowheads in H&E images indicate cells with fusiform morphology, indicative of osteoblasts. White arrowheads in osteopontin images indicate areas adjacent to scaffold-derived hydroxyapatite. Detailed inset shown for TRAP to highlight multi-nucleated TRAP+ cells (Scale bar: 50 μm).

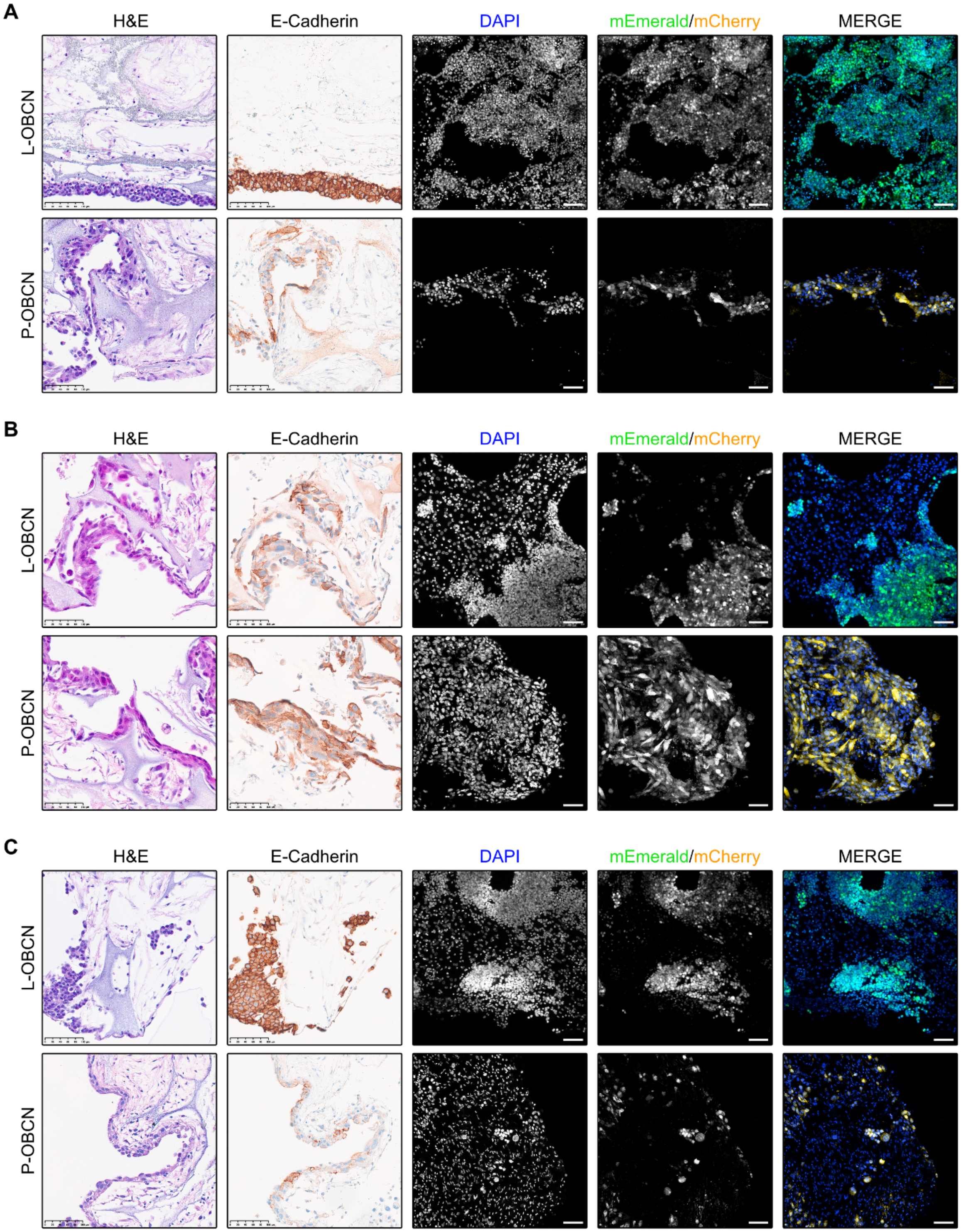

**Supplementary Figure 3 - Histological and immunofluorescence analysis of LNCaP and PC3 cell engraftment in OBCNs.** Representative images from L-OBCN and P-OBCN niches derived from: (A) donor BM256 at TP1, (B) BM267 at TP2, and (C) BM256

at TP2. Samples were analyzed by H&E staining, E-Cadherin IHC, and whole-mount IF. Fluorescent channels include DAPI (nuclei), mEmerald (LNCaP), and mCherry (PC3). Individual channels and merged images are shown. Scale bars: 100 µm.

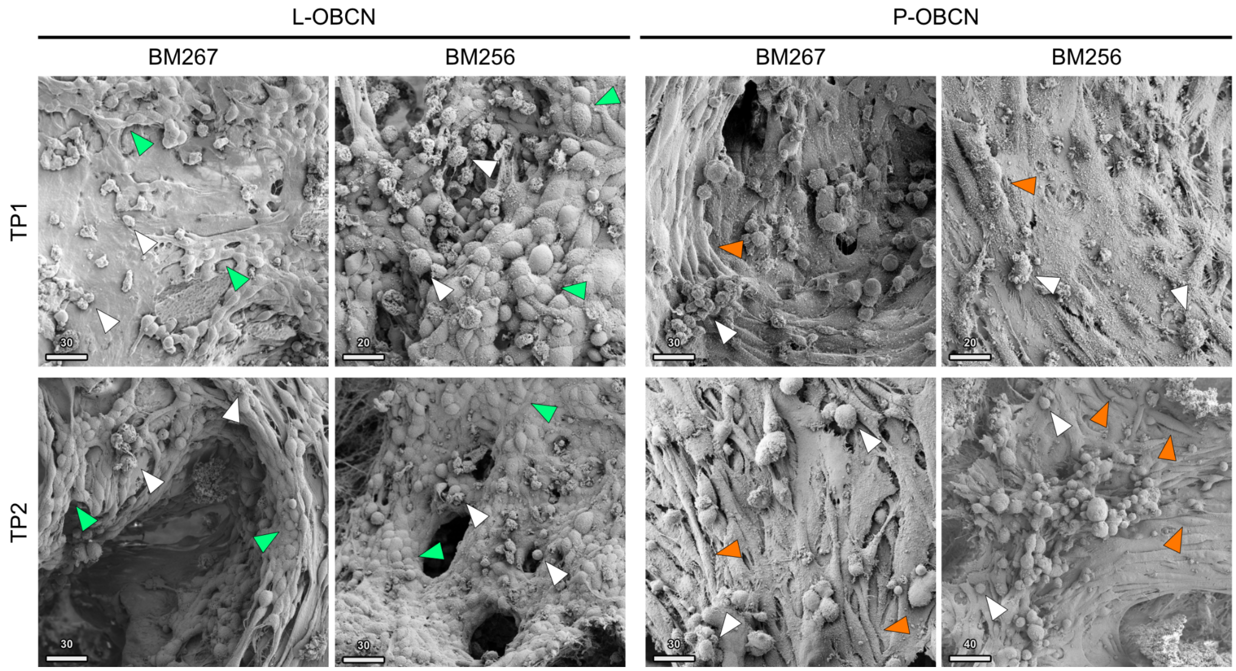

**Supplementary Figure 4 - Scanning electron microscopy (SEM) of prostate cancer cell engraftment in the niche.** SEM images of L-OBCNs and P-OBCNs generated from donors BM267 and BM256, acquired at day 4 (TP1) and day 10 (TP2) of co-culture with PCa cells. LNCaP and PC3 cells are indicated by green and orange arrowheads, respectively. White arrowheads highlight monocyte-derived cells within the niche. Scale bars: 20–40 µm.

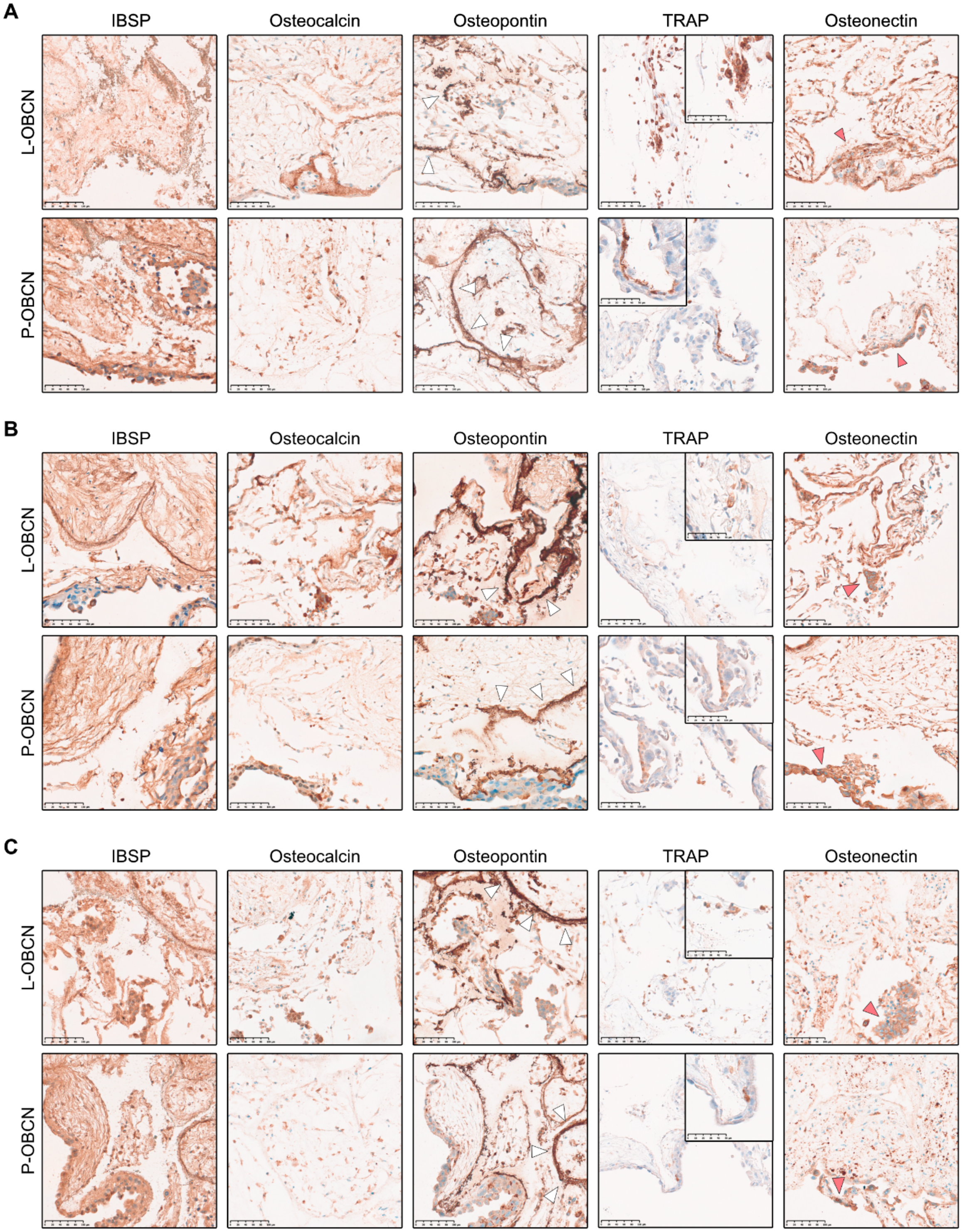
A
IBSP
Osteocalcin
Osteopontin
TRAP
Osteonectin
L-OBCN
P-OBCN
B
IBSP
Osteocalcin
Osteopontin
TRAP
Osteonectin
L-OBCN
P-OBCN
C
IBSP
Osteocalcin
Osteopontin
TRAP
Osteonectin
L-OBCN
P-OBCN

**Supplementary Figure 5 - Histological assessment of bone-related markers in PCa co-cultured OBCNs.** Representative IHC staining for key bone microenvironment markers (IBSP, osteocalcin, osteopontin, TRAP, and osteonectin) in L-OBCNs and P-OBCNs derived from: (A) BM256 at TP1, (B) BM267 at TP2, and (C) BM256 at TP2. White arrowheads in osteopontin panels highlight areas adjacent to scaffold-derived hydroxyapatite. Red arrowheads indicate PCa cells positive for osteonectin. High-magnification insets in TRAP panels show multinucleated $TRAP^+$ cells. Scale bars: 100 µm (main panels), 50 µm (TRAP insets).

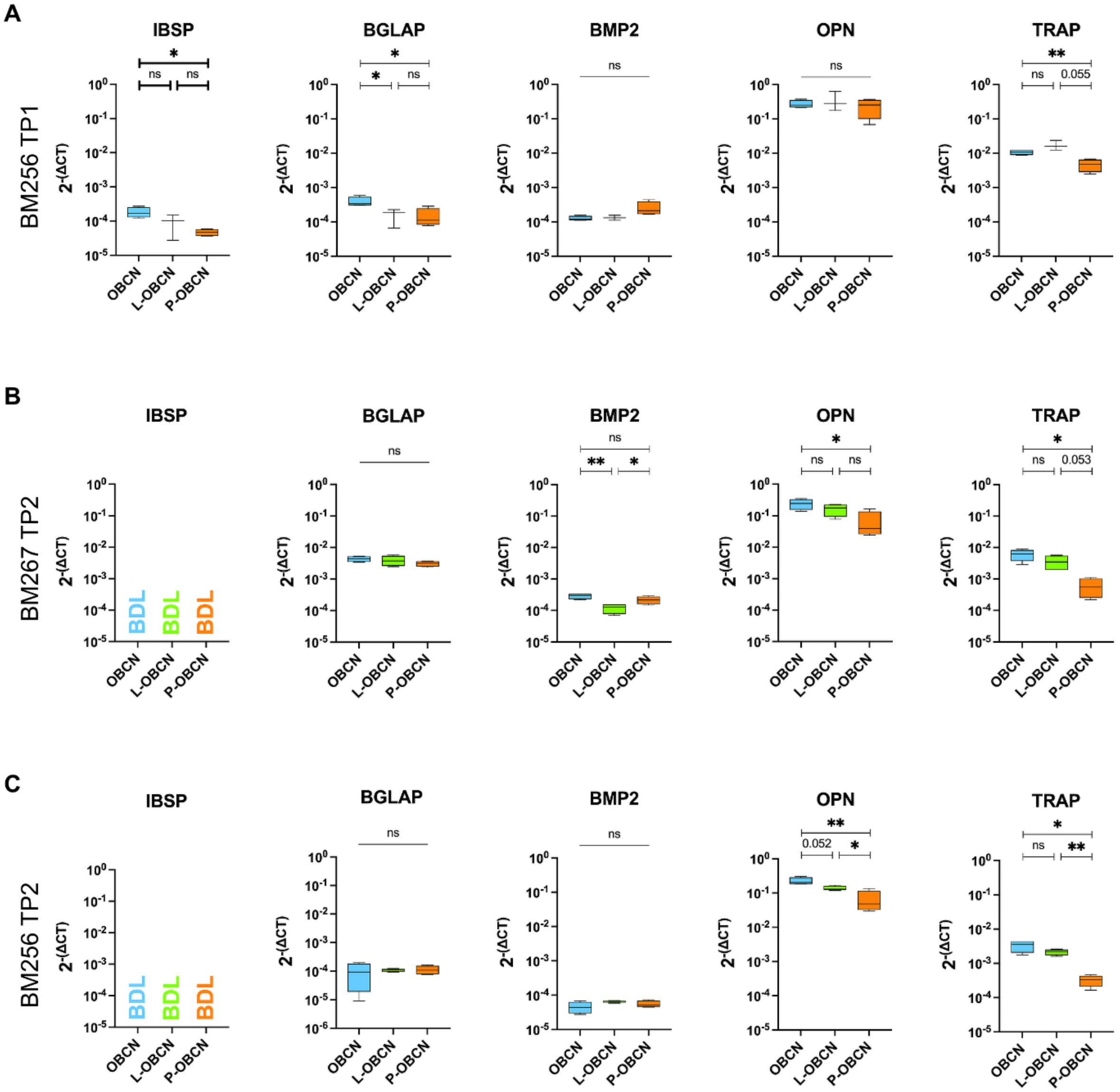

**Supplementary Figure 6 - Gene expression analysis of key bone-related markers in non-tumor niche cells sorted from OBCNs, L-OBCNs, and P-OBCNs.** qRT-PCR analysis of osteoblastic and osteoclastic markers IBSP, BGLAP, BMP2, OPN, and TRAP on osteoblasts and osteoclasts retrieved from the OBCNs after sorting, for (A) donor BM256 at TP1, (B) BM267 at TP2, and (C) BM256 at TP2. Comparisons are drawn between cells extracted from OBCNs without PCa (blue), L-OBCNs (green), and P-OBCNs (orange). Comparisons were done via Welch t-test, with statistically significant changes being shown in the graph (∗: $p < 0.05$; ∗∗: $p < 0.005$; ∗∗∗: $p < 0.0005$) with n=4 experimental replicates. Samples with undetectable gene expression have been marked with BDL (below detection level).

A

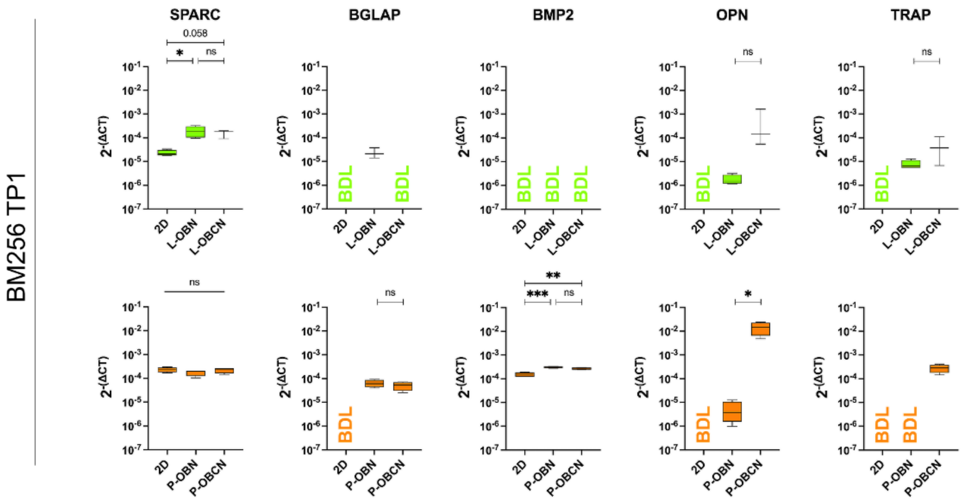
BM256 TP1
SPARC
BGLAP
BMP2
OPN
TRAP
0.058
*
ns
**
***
2-(ΔCT)
BDL
2D
L-OBN
L-OBCN
P-OBN
P-OBCN

B

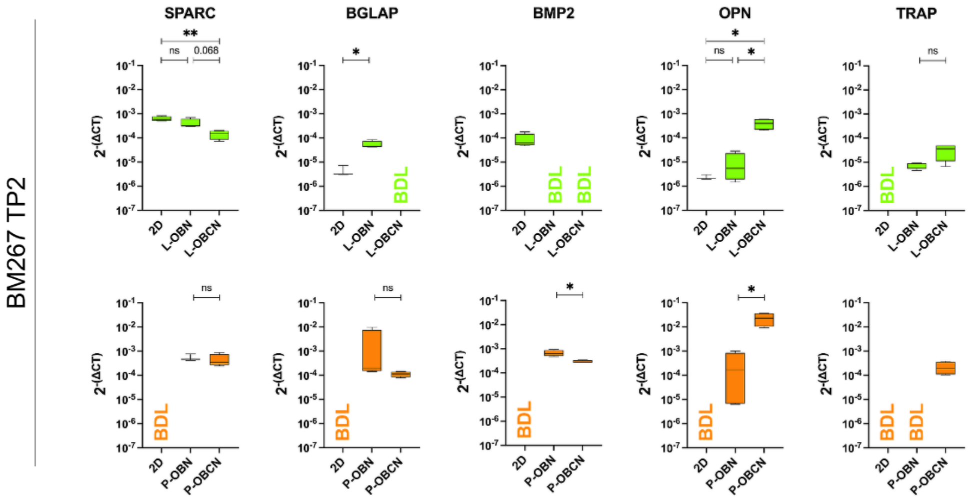
BM267 TP2
SPARC
BGLAP
BMP2
OPN
TRAP
**
ns
0.068
*
2-(ΔCT)
BDL
2D
L-OBN
L-OBCN
P-OBN
P-OBCN

C

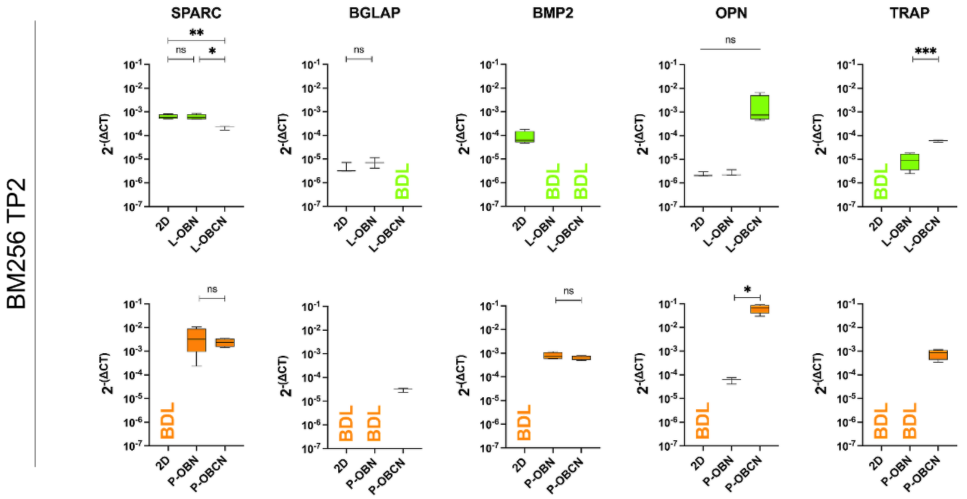
BM256 TP2
SPARC
BGLAP
BMP2
OPN
TRAP
**
ns
*
***
2-(ΔCT)
BDL
2D
L-OBN
L-OBCN
P-OBN
P-OBCN

**Supplementary Figure 7 - Analysis of gene expression of bone-related markers in PCa cell lines extracted from L-OBCNs and P-OBCNs compared to their 2D-grown counterparts.** qRT-PCR analysis of osteoblastic and osteoclastic markers SPARC, BGLAP, BMP2, OPN, and TRAP on PCa cells extracted from L-OBNs and L-OBCNs (green) or P-OBNs and P-OBCNs (orange) for (A) BM256 at TP1, (B) BM267 at TP2, and (C) BM256 at TP2. Comparisons are drawn between cells co-cultured in the OBCNs and the same kind of PCa line grown in 2D under the same medium conditions. Comparisons were done via Welch t-test, with statistically significant changes being shown in the graph (✱: $p < 0.05$; ✱✱: $p < 0.005$; ✱✱✱: $p < 0.0005$) with n=4 experimental replicates. Samples with undetectable gene expression have been marked with BDL (below detection level).

### 3. Reagents and Instruments List

The section number for each table refers to the section in the manuscript.

| 2.1 Cell Culture Media | | | |
|---|---|---|---|
| **Reagent / Component Name** | **Supplier** | **Catalog Number** | **Concentration** |
| Minimum Essential Medium Alpha (aMEM) | Gibco | 22571 | Pure |
| Fetal Bovine Serum (FBS) | Gibco | A5256701 | 10% |
| GlutaMAX | Gibco | 35050 | 1% |
| Penicillin-Streptomycin | Gibco | 15140 | 1% |
| HEPES | Gibco | 15630 | 1% |
| Sodium Pyruvate | Gibco | 11360 | 1% |
| Fibroblast Growth Factor 2 (FGF2) | R&D Systems | 233-FB | 5 ng/mL |
| Ascorbic Acid | Sigma-Aldrich | A8960 | 0.1 mM |
| Dexamethasone | Sigma-Aldrich | D2915 | 10 nM |
| Beta-Glycerophosphate | Sigma-Aldrich | G9422 | 10 mM |
| 1,25-Dihydroxyvitamin D3 | Tocris | 2551 | 10 nM |
| Macrophage Colony-Stimulating Factor (M-CSF) | Gibco | AF-300-25 | 25 ng/mL |

| 2.2 Cell Source and Cell Expansion | | |
|---|---|---|
| **Reagent / Component Name** | **Supplier** | **Catalog Number** |
| Ficoll-Paque PLUS | Cytiva | 17144002 |
| CD14 MicroBeads | Miltenyi Biotec | 130-050-201 |
| LS Columns | Miltenyi Biotec | 130-042 |
| QuadroMACS Separator | Miltenyi Biotec | 130-091-051 |
| **Cell Line Name** | **Supplier** | **Catalog Number** |
| LNCaP prostate cancer cell line | DMSZ | ACC256 |
| PC3 prostate cancer cell line | ATCC | CRL-1435 |

| **Prostate organoid medium (POM) components:** | **Supplier** | **Catalog Number** | **Concentration** |
|---|---|---|---|
| AdDMEM-F12 | Gibco | 12634010 | Pure |
| Glutamax | Gibco | 35050061 | 1% |
| Hepes | Gibco | 15630056 | 1% |
| Pen/Strep | BioConcept | 4-01F00-H | 1% |
| Primocin | InvivoGen | ant-pm-1 | 100 µg/ml |
| B27 | Life Technologies | 17504044 | 1X |
| Nicotinamide | Sigma-Aldrich | N0636-100g | 10 mm |
| N-Acetyl-l-cysteine | Sigma-Aldrich | A9165-25g | 1.25 mm |
| A83-01 | Tocris | 2939 | 500 nm |
| FGF-10 | Peprotech | 100-26-25UG | 10 ng/ml |
| FGF-2 | Peprotech | 100-18B-50UG | 5 ng/ml |
| EGF | Peprotech | AF-100-15-100UG | 5 ng/ml |
| Noggin | Peprotech | 120-10C-50UG | 100 or 50 ng/ml |
| Rspondin 1 | R&D | 4645-RS-100 | 500 or 250 ng/ml |
| Prostaglandin E2 | Tocris | 2296 | 1 µm |
| SB202190 | Sigma-Aldrich | S7076-5MG | 10 µm |
| Y-27632 dihydrochloride | Abmole Bioscience | M1817 | 10 µm |
| (Dihydro)testosterone (5α-androstan-17β-ol-3-one) | Sigma-Aldrich | 10300-5G-F | 1 or 0.1 nm |

143

| **2.3 3D Niche Co-culture** | | |
|---|---|---|
| **Reagent / Component Name** | **Supplier** | **Catalog Number** |
| Porous hydroxyapatite scaffold (8×4 mm) | Finceramica | ENGI-PROD-01 |
| Low-attachment 12-well plates | Sarstedt | 4021721 |

144

| **2.4 Cell Isolation from the 3D Constructs** | | | |
|---|---|---|---|
| **Reagent / Component Name** | **Supplier** | **Catalog Number** | **Concentration** |
| Type II Collagenase | Worthington | LS004176 | 0.15% |

| | | | |
|---|---|---|---|
| Thermomixer C | Eppendorf | 5382000015 | — |
| Trypsin-EDTA (0.25%) | Gibco | 25200072 | — |
| 70 µm Cell Strainers | Corning | 352350 | — |

| **2.5 Sorting of mEmerald-LNCaP and mCherry-OC3 cells from OBNs and OBCNs** | | |
|---|---|---|
| **Reagent / Component Name** | **Supplier** | **Catalog Number** |
| CytoFLEX SRT | Beckman Coulter | |
| FACSAria III | BD Biosciences | |

| **2.6 RNA extraction, cDNA synthesis, and qRT-PCR analysis** | | |
|---|---|---|
| **Reagent / Equipment Name** | **Supplier** | **Catalog Number** |
| RNeasy Mini Kit | Qiagen | 74106 |
| RNeasy Micro Kit | Qiagen | 74004 |
| NanoDrop One Spectrophotometer | Thermo Fisher | ND-ONE-W |
| SuperScript III Reverse Transcriptase | Thermo Fisher | 18080093 |
| ViiA 7 Real-Time PCR System | Thermo Fisher | — |
| TaqMan Assay – IBSP | Thermo Fisher | Hs00173720_m1 |
| TaqMan Assay – BGLAP | Thermo Fisher | Hs01587814_g1 |
| TaqMan Assay – BMP2 | Thermo Fisher | Hs00154192_m1 |
| TaqMan Assay – ALPL | Thermo Fisher | Hs01029144_m1 |
| TaqMan Assay – OPN | Thermo Fisher | Hs00959010_m1 |
| TaqMan Assay – TRAP | Thermo Fisher | Hs00356261_m1 |
| TaqMan Assay – SPARC | Thermo Fisher | Hs00234160_m1 |
| TaqMan Assay – GAPDH (endogenous control) | Thermo Fisher | Hs02758991_g1 |

| **2.7 Whole-mount immunofluorescence staining and confocal microscopy** | | | |
|---|---|---|---|
| **Reagent / Equipment Name** | **Supplier** | **Catalog Number** | **Concentrat** |

| | | | ion |
|---|---|---|---|
| Formaldehyde 4% | Thermo Fisher | 28908 | 4% |
| Triton X-100 | Sigma-Aldrich | 93443 | 0.40% |
| Bovine Serum Albumin (BSA) | Sigma-Aldrich | A9647 | 2% |
| Goat Serum | Gibco | 16210064 | 1%, 5%, or 10% |
| Anti-Cytokeratin-8 (CK8) Antibody | Biolegend | 904804 | 1 : 200 |
| Anti-TRAP Antibody | Abcam | AB185716 | 1 : 200 |
| Goat Anti-Mouse 546 (Secondary) | Invitrogen | A11030 | 1 : 200 |
| Goat Anti-Rabbit 647 (Secondary) | Invitrogen | A21244 | 1 : 200 |
| DAPI | BD Biosciences | 564907 | 1 mg/mL |
| Imaging Dishes (35 mm) | Ibidi | 81218-200 | — |
| Ti2-E Inverted Microscope | Nikon | — | — |
| AxR Point-Scanning Confocal Unit | Nikon | — | — |
| NIS Elements Software | Nikon | — | — |

| **2.8 Histological processing, immunohistochemistry, and immunofluorescence staining** | | | |
|---|---|---|---|
| **Reagent / Equipment Name** | **Supplier** | **Catalog Number** | **Concentration** |
| HistoGel | Epredia | HG-4000-012 | Pure |
| EDTA | Sigma-Aldrich | E6635 | 15% |
| TPC-15 Trio | MEDITE Medical | — | — |
| Microtome | Thermo Scientific | — | — |
| SuperFrost Plus Glass Slides | Fisherbrand | — | — |
| Xylene (Xylol Histograde) | Biosystems | 3410 | Pure |
| Citrate Buffer pH 6 (50x) | quartett Q Retrieval | AR-001-0120 | 1 : 50 |
| Triton X-100 | Sigma-Aldrich | 93443 | 0.40% |
| Goat Serum | Gibco | 16210064 | 1% |
| Anti-Cytokeratin-8 (CK8) Antibody | Biolegend | 904804 | 1 : 200 |
| Anti-Osteocalcin (OCN) Antibody | Thermo Fisher | PA5-96529 | 1 : 200 |
| Anti-E-Cadherin (ECAD) Antibody | Ventana | 760-4440 | ready-to-use |

| | | | |
|---|---|---|---|
| Anti-TRAP Antibody | Bio BS | BSB5982 | ready-to-use |
| Anti-Bone Sialoprotein (BSP) | Abcam | AB52128 | 1 : 100 |
| Anti-Osteonectin (ON) | Thermo Fisher | 33-5500 | 1 : 500 |
| Anti-Osteopontin Antibody (OCN) | Proteintech | 22952-1-AP | 1 : 200 |
| Goat Anti-Rabbit 488 (Secondary) | Invitrogen | A11034 | 1 : 200 |
| Goat Anti-Mouse 647 (Secondary) | Invitrogen | A21236 | 1 : 200 |
| DAPI | BD Biosciences | 564907 | 1 mg/mL |
| Ventana BenchMark ULTRA | Roche Diagnostics | — | — |
| Fluoromount Aqueous Mounting Medium | Sigma-Aldrich | F4680 | — |
| Gemini AS Automated Stainer | Epredia | — | — |
| NanoZoomer S60 Slide Scanner | Hamamatsu | — | — |
| Ti2 Widefield Fluorescence Microscope | Nikon | — | — |
| DS-Ri2 Camera | Nikon | — | — |
| NIS Elements AR Software (v5.21.03) | Nikon | — | — |

| **2.9 Quantification of PCa Organoids Aggregates** | | |
|---|---|---|
| **Reagent / Software / Equipment Name** | **Supplier** | **Version / Catalog Number** |
| Anti-Cytokeratin-8 (CK8) Antibody | Biolegend | 904804 |
| DAPI | BD Biosciences | 564907 |
| Nikon AxR Confocal Microscope | Nikon | — |
| Imaris File Converter | Bitplane | v10.1.10 |
| Imaris with Measurement Pro Module | Bitplane | v10.1.10 |

| **2.10 Statistics** | | |
|---|---|---|
| **Software / Method Name** | **Supplier** | **Version / Details** |
| GraphPad Prism | GraphPad | v10.4.2 |

151

| 9.1 Lentiviral Labelling of PCa Cell Lines | | | |
|---|---|---|---|
| **Reagent / Component Name** | **Supplier** | **Catalog Number** | **Concentration** |
| pLVX lentiviral vector backbone | Clontech/Takara Bio | 632164 | |
| Lenti-X 293T packaging cells | Clontech/Takara Bio | 632180 | |
| Packaging plasmid psPAX2 | Addgene | 12260 | |
| Packaging plasmid pMD2.G | Addgene | 12259 | |
| Packaging plasmid pRSV-Rev (likely) | Addgene | 12253 | |
| Lipofectamine 2000 | Thermo Fisher | 11668027 | |
| Polybrene | Sigma-Aldrich | H9268 | 8 μg/mL |
| G418 (Geneticin) | Roche Diagnostics | 108321-42-2 | 250 μg/mL |

152

| 9.5 Scanning Electron Microscopy Imaging | | | |
|---|---|---|---|
| **Reagent / Equipment Name** | **Supplier** | **Catalog Number** | **Concentration** |
| Glutaraldehyde | Thermo Scientific | A17876 | 2% |
| Sodium Cacodylate Buffer | Thermo Scientific | J62202.AE | 0.1M |
| Critical Point Dryer | Tousimis | Autosamdri-815 | — |
| Sputter Coater | LEICA | EM ACE600 | — |
| SEM Microscope | ZEISS | Gemini 2 | — |

153
